# IκBα loss impairs mitochondrial homeostasis and primes cells for senescence

**DOI:** 10.64898/2026.11.20.767802

**Authors:** Keita Matsuda, Takuma Kono, Aeryeong Hyun, Kayo Hibino, Shou Soeda, Sho Tabata, Shashank Tiwari, Björn Goldenbogen, Masatomo Takahashi, Yoshihiro Izumi, Takeshi Bamba, Jun Adachi, Helfrid Hochegger, Jana Wolf, Mariko Okada

## Abstract

Inflammatory signaling and mitochondrial dysfunction are hallmarks of cellular senescence, yet how these processes are coordinated remains incompletely understood. Here, we identify IκBα, a well-known inhibitor of NF-κB, as a regulatory factor that links mitochondrial homeostasis to stress-induced cell death and senescence. Knockdown (KD) of IκBα did not by itself induce a fully established senescent state but instead generated a senescence-primed state characterized by cellular hypertrophy, reduced DNA synthesis, metabolic remodeling, and mitochondrial dysfunction. Time-course proteomic and phosphoproteomic analyses revealed a selective reduction of mitochondrial inner membrane-associated proteins in IκBα KD cells and attenuated early signaling responses to TNFα stimulation, respectively. Single-cell transcriptomic analysis further indicated that a smaller fraction of IκBα KD cells was predicted to transition toward a proapoptotic state. Among canonical IκB family members, induction of the senescence-associated secretory phenotype (SASP) was specific to IκBα KD, even though mitochondrial dysfunction was more pronounced in IκBβ and IκBε KD. These findings suggest that IκBα influences the balance between post-stress cellular senescence and cell death through mechanisms that include maintaining mitochondrial homeostasis.

## INTRODUCTION

The NF-κB signaling pathway plays an important role in regulating cell function and fate in a variety of cell types, and its aberrant regulation contributes to many human diseases^1,2^. The pathway is activated by stimulation of several classes of receptors, including innate immune receptors, antigen receptors, and inflammatory cytokine receptors such as the tumor necrosis factor (TNF) receptor. Receptor activation triggers a signaling cascade that activates IκB kinase β (IKKβ), which phosphorylates the inhibitory κB (IκB) proteins (IκBα, IκBβ, and IκBε) and targets them for proteasomal degradation^3,4^. Because IκBα binds NF-κB and masks the nuclear localization signal of RelA (p65), its phosphorylation-dependent degradation allows the p50/RelA (p65) heterodimer to translocate into the nucleus^5^. Among the three IκB proteins, IκBα (NFKBIA) is a well-known negative feedback regulator that limits the duration of NF-κB activity^5–7^.

Cellular senescence is a stable state of cell-cycle arrest induced by DNA damage, oxidative stress, replicative stress, and other environmental factors^8–10^. Senescent cells exhibit diverse phenotypes, including cell enlargement, mitochondrial dysfunction, and metabolic rewiring^11–14^, as well as the senescence-associated secretory phenotype (SASP), which is characterized by the secretion of inflammatory cytokines and chemokines that influence neighboring cells and the tissue microenvironment^15,16^. NF-κB acts as a transcription factor that induces target genes, including those that constitute the SASP, and thereby contributes to the establishment and maintenance of cellular senescence^17–20^. In our previous study, loss of IκBα sustained NF-κB nuclear localization after TNFα stimulation, enhanced its DNA-binding activity, and increased the expression of NF-κB target genes, including SASP-related factors^21^. These findings suggest that IκBα loss can promote a senescence-associated inflammatory state through sustained NF-κB activity. Although these senescence-associated phenotypes often occur together, how they are mutually regulated and how they contribute to the establishment and maintenance of senescence remain unclear.

Mitochondria are central organelles for ATP production, redox homeostasis, and metabolic function, and impaired mitochondrial homeostasis, including reduced respiration and altered membrane potential, has been reported in senescent cells^22–24^. Mitochondrial dysfunction has also been shown to contribute to SASP induction^11,25^, and it may act as a trigger of senescence rather than merely a consequence of it^25–27^. These findings imply a close relationship between mitochondrial dysfunction and inflammatory transcriptional responses, raising the possibility that NF-κB links the two during cellular senescence. In this context, IκBα, the major inhibitor of NF-κB, is of particular interest, because it has also been reported to localize to mitochondria and suppress cell death^28–31^, suggesting roles in mitochondrial regulation and cell fate beyond the control of NF-κB-dependent transcription.

Here, we investigated whether IκBα regulates mitochondrial function and senescence, independently of or in addition to its role in NF-κB signaling. We found that IκBα knockdown (KD) impaired mitochondrial homeostasis even without stimulation, accompanied by a selective loss of mitochondrial inner membrane-associated proteins and altered cellular phenotypes.

Phosphoproteomic and metabolomic analyses linked the reduced ATP levels in IκBα KD cells to attenuated early phosphorylation responses to TNFα. Single-cell transcriptomic analysis further revealed that IκBα KD shifted the balance of cell-state transitions after TNFα stimulation, away from a pro-apoptotic state and toward an anti-apoptotic, senescence-associated state.

Comparison with IκBβ and IκBε KD showed that SASP induction was specific to IκBα KD, even though mitochondrial dysfunction was more pronounced in IκBβ and IκBε KD. These results suggest that impaired mitochondrial homeostasis may contribute to the senescence-prone state of IκBα KD cells. As IκBα reduction also enhances TNFα-induced, NF-κB-dependent inflammatory transcription in the nucleus, this nuclear function and the mitochondrial changes may cooperate to promote senescence-associated phenotypes.

## RESULTS

### IκBα reduction promotes stress-induced senescence while limiting cell death

We previously showed that IκBα reduction promotes senescence under TNFα stimulation^21^. To test whether this effect extends to other types of stimuli, we used human breast cancer MCF-7 cells, which are relatively resistant to replicative senescence^32,33^, and measured senescence-associated β-galactosidase (SA-β-gal) positivity after exposure to inflammatory (TNFα), oxidative (H_2_O_2_), or DNA damage stress (etoposide or camptothecin) for 24 or 48 h (Fig. 1A-D). IκBα knockdown (KD) alone did not markedly increase the fraction of SA-β-gal-positive cells (median, control, 9.8%; IκBα KD, 12.0%; P = 0.170), whereas under each stress condition, the fraction of SA-β-gal-positive cells was significantly higher in IκBα KD cells than in control cells at 24 and 48 h (P < 0.05). These results indicate that IκBα reduction sensitizes cells to stressinduced cellular senescence.

**Figure 1.**
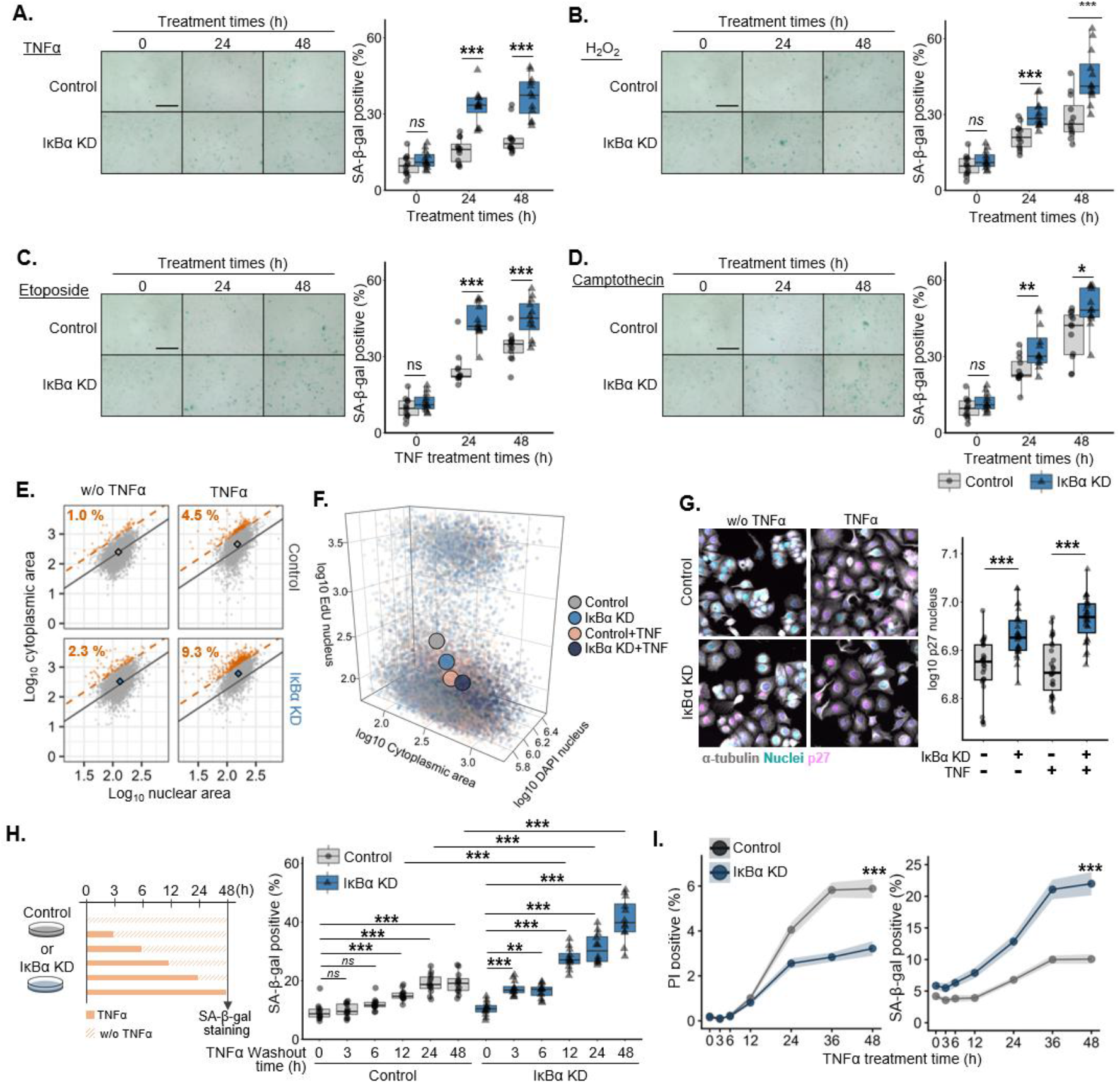
IκBα depletion enhances stress-induced senescence and limits TNFα-induced cell death MCF-7 cells were transfected with control or IκBα siRNA (25 nM). (A–D) SA-β-gal positivity after treatment with TNFα (10 ng/mL) (A), H₂O₂ (300 μM) (B), etoposide (10 μM) (C), or camptothecin (10 μM) (D) for 24 or 48 h. Representative images and quantification are shown. Scale bars, 100 μm. (E) Single-cell analysis of nuclear and cytoplasmic areas before stimulation or after TNFα treatment (10 ng/mL, 24 h). Robust regression in unstimulated Control cells at 0 h defined the reference relationship. Cells with residuals above the 99th percentile of the reference distribution were classified as cytoplasmically enlarged (orange; dashed line, threshold). Diamonds indicate condition medians. (F) Three-dimensional single-cell analysis of cytoplasmic area, nuclear DAPI content, and EdU signal before stimulation or after TNFα treatment (10 ng/mL, 24 h). All variables were log₁₀-transformed. Large points indicate the mean value along each axis for each condition. (G) Nuclear p27 intensity in Control and IκBα KD cells with or without TNFα treatment (10 ng/mL, 24 h), shown on a log₁₀ scale. (H) Effect of TNFα exposure duration on SA-β-gal positivity. Left: Cells were exposed to TNFα (10 ng/mL) for 0, 3, 6, 12, 24, or 48 h, followed by washout and culture in TNFα-free medium until 48 h. Orange and hatched bars indicate TNFα treatment and post-washout culture, respectively. Right: SA-β-gal positivity after the indicated exposure durations. (I) Time-course of SA-β-gal positivity (left) and PI positivity (right) after TNFα treatment (10 ng/mL). Points indicate means and shaded areas indicate SE. For visualization in (E) and (F), up to 5,000 cells were randomly sampled per condition; in (F), each axis was restricted to the 1st–99th percentile range. Box plots in (G) and (H) show the median and interquartile range, with whiskers extending to 1.5 × the interquartile range. For (A)–(D) and (H)–(I), n = 12 fields of view per group; for (G), n = 27 fields of view per group. P values in (A)–(D), (G), and (I) were determined using unpaired two-tailed Welch’s t-tests for the indicated comparisons. In (H), comparisons with 0 h within each siRNA condition were performed using Dunnett’s multiple comparisons test, whereas comparisons between Control and IκBα KD at 12, 24, and 48 h were performed using unpaired two-tailed Welch’s t-tests with Holm correction. *P < 0.05, **P < 0.01, ***P < 0.001.

Next, we analyzed cell morphology and DNA synthesis (EdU incorporation) in control and IκBα KD cells, with or without TNFα stimulation (24 h), using elution-based iterative indirect immunofluorescence (4i)^34^ (see Methods). IκBα KD increased the proportion of enlarged cells even without stimulation (control, 1.0%; IκBα KD, 2.3%), and this increase was further enhanced by TNFα stimulation (control, 4.5%; IκBα KD, 9.3%) (Fig. 1E). Single-cell analysis of cytoplasmic area, nuclear DAPI intensity, and EdU incorporation further revealed a marked shift in cellular state from untreated control cells to TNFα-treated IκBα KD cells, reflecting increased cell size and reduced DNA synthesis (Fig. 1F). These results suggest that IκBα KD promotes a senescence-prone state even in the absence of external stress.

We further examined p27 (CDKN1B) expression using immunofluorescence. We chose p27 because MCF-7 cells lack p16INK4A (CDKN2A)^35,^ and no increase in p53 or p21 levels was observed in our 4i analysis (Fig. S1A and S1B). p27 intensity was higher in IκBα KD cells than in control cells both without stimulation (+16.8%, P = 2.8 × 10⁻⁵) and after 24 h of TNFα stimulation (+26.3%, P = 2.7 × 10⁻⁹) (Fig. 1G). However, although increased p27 is associated with cell-cycle inhibition, it is also observed in reversible growth arrest^36,37^. We therefore tested whether the TNFα-induced senescence-associated phenotypes were transient or persisted after stimulus removal. Cells were treated with TNFα for defined periods, followed by incubation in TNFα-free medium for the remaining time to complete a total of 48 h before SA-β-gal-positivity was assessed (Fig. 1H). After TNFα treatment and washout, the fraction of SA-β-gal-positive cells increased from 9.4% (0 h) to 15.1% (12 h) in control cells and from 10.5% to 27.6% in IκBα KD cells. At 12, 24, and 48 h, the fraction was higher in IκBα KD cells than in control cells (27.6% vs. 15.1%, 31.0% vs. 19.1%, and 40.8% vs. 19.2%, respectively; all adjusted P < 0.001) (Fig. 1H). These results suggest that IκBα KD promotes a persistent, senescence-associated state after transient TNFα exposure.

Because senescent cells often depend on anti-apoptotic pathways and resist apoptotic stimuli ^38,39^, we next examined whether IκBα KD alters cell death after TNFα stimulation, using propidium iodide (PI) staining. TNFα (48 h) increased the fraction of PI-positive cells in control cells (5.88%), whereas this increase was suppressed by IκBα KD (3.22%) (P < 0.001) (Fig. 1I). This trend was opposite to the increase in SA-β-gal-positive cells under IκBα KD, which reached 21.99% compared with 10.07% in control cells at 48 h (P < 0.001). Together with the results above, these findings suggest that IκBα reduction alone does not fully induce cellular senescence, but promotes a senescence-prone state in which cells survive TNFα stimulation and are likely to acquire senescence-associated phenotypes.

### Early stress-adaptive kinase responses to TNFα are blunted in IκBα KD cells

To comprehensively evaluate the effect of IκBα reduction on senescence, we acquired time-series transcriptomic, proteomic, and phosphoproteomic data after TNFα stimulation, covering 14,276 genes, 8,588 proteins, and 13,170 phosphosites, respectively (Fig. S2A; see Methods for details). To assess the overall responses to TNFα, we used each condition’s 0-h samples as its baseline and counted the phosphosites, genes, and protein groups that changed at each post-stimulation time point (BH-adjusted P < 0.05 and |response| ≥ 0.5). Response was defined as the centered phosphosite value difference, the RNA log₂ fold change, and the protein log₂ abundance difference, respectively. IκBα KD reduced the number of changed features most strongly in the phosphoproteome, both in total across time points (−75.1% vs. control) and in unique features, defined as distinct features that changed significantly at one or more time points (−69.7%) (Fig. 2A). The transcriptome was moderately affected (total, −12.4%; unique,

**Figure 2.**
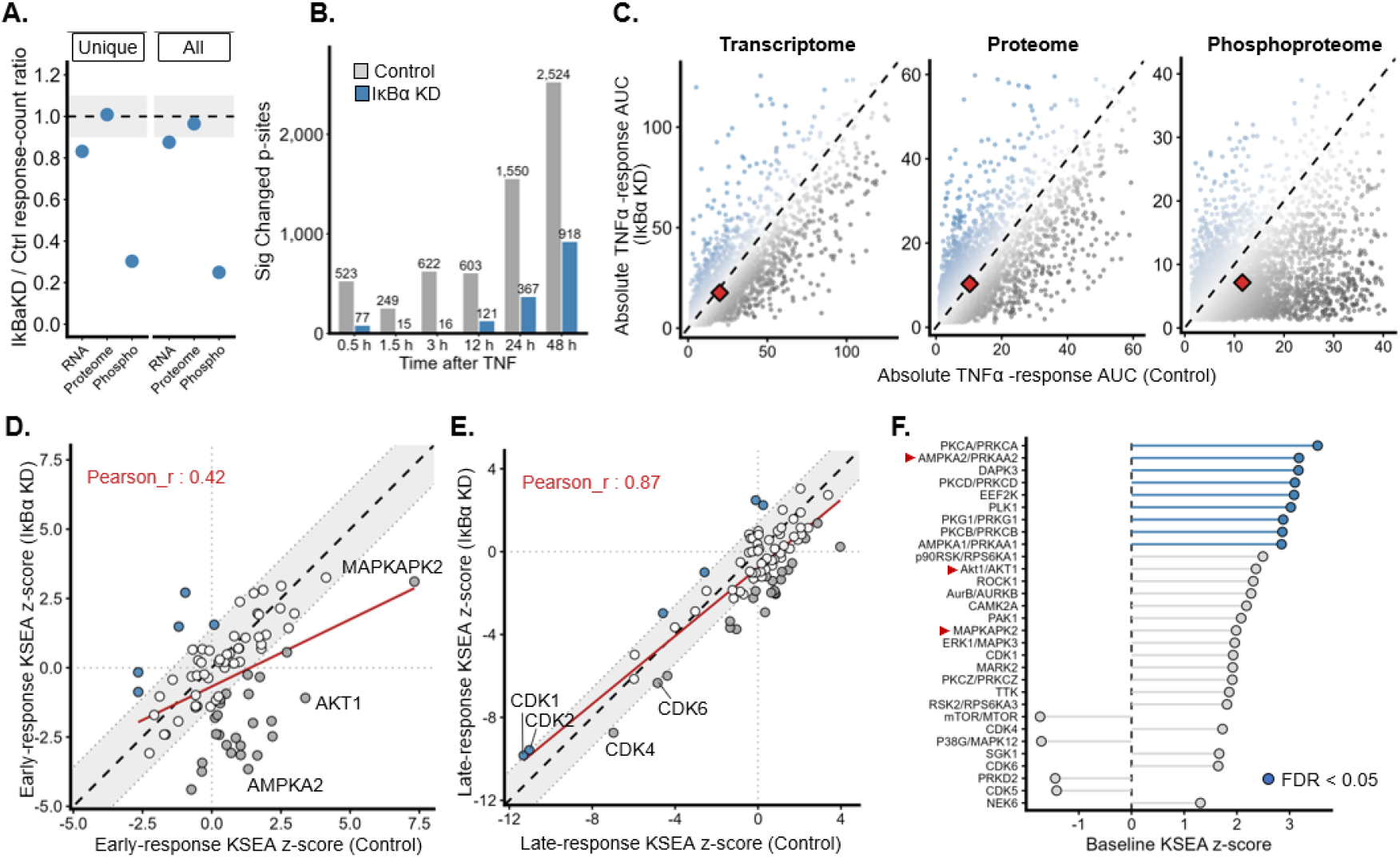
IκBα depletion preferentially blunts early stress-adaptive kinase responses to TNFα (A) Ratio of TNF-responsive features in IκBα KD versus Control across transcriptome, proteome, and phosphoproteome layers. Unique responders were features significant at ≥1 time point, whereas feature-time burden represents the total number of significant feature–time point pairs. Dashed line, ratio = 1; shading, 0.9–1.1. (B) Number of TNF-responsive phosphosites at each time point in Control and IκBα KD cells. (C) Absolute AUC of TNF responses in Control versus IκBα KD for features with complete 0–48 h trajectories. AUCs were calculated by trapezoidal integration of absolute responses, with the 0-h response set to 0. Dashed line, y = x; diamond, centroid. Up to 6,000 features per layer were randomly sampled and displayed within the 1st–99th percentile range. (D, E) Kinase activity responses in Control and IκBα KD cells during the early (0–3 h; D) and late (12–48 h; E) TNF-response windows. Time-averaged signed phosphosite responses were used for KSEA. All analyzed kinases with ≥5 mapped phosphosites are shown. Gray region, perpendicular distance of ±1 from y = x; red line, linear regression. Pearson’s r indicates the correlation between Control and IκBα KD KSEA z-scores. (F) Baseline KSEA comparing IκBα KD and Control cells at 0 h. KSEA was calculated from phosphosite-level IκBα KD–Control differences, and the 30 kinases with the largest absolute KSEA z-scores are shown. Positive and negative z-scores indicate IκBα KD-high and Controlhigh activity, respectively. For (A)–(E), TNF responses were calculated relative to the respective 0-h condition. Significant TNF responses in (A) and (B) were defined as BH FDR < 0.05. and absolute response ≥ 0.5. For (D)–(F), n = 3 biological replicates per condition at each analyzed time point. Significant kinases in (F) were defined as BH FDR < 0.05.

−16.8%), whereas the proteome was largely unchanged (total, −3.5%; unique, +0.9%). We next examined the numbers of significantly changed features at individual time points. In the phosphoproteome, the number of significantly changed phosphosites was lower in IκBα KD cells by more than 100 at every time point examined (Fig. 2B). In the transcriptome, the reduction was most pronounced at 48 h after stimulation (−1,132 genes; −29.2%) (Fig. S2B), whereas the reduction across all time points was more moderate. In contrast, the proteome showed differences of fewer than 100 significantly changed protein groups between control and IκBα KD cells at each time point (Fig. S2C).

To distinguish whether this reflected a threshold effect of the analysis or attenuation of signal response in the cells, we calculated dynamic indices for each phosphorylation site relative to the pre-stimulation state (AUC, maximum peak, dynamic range, and maximum slope; see Methods) and compared them between control and IκBα KD cells (Fig. 2C and Fig. S2D). In the phosphoproteome, most phosphorylation sites were distributed below the identity line (y = x), indicating that phosphorylation changes induced after TNFα stimulation were globally reduced under IκBα KD (71.0% of phosphosites showed a greater absolute AUC in Control than in IκBα KD cells; Fig. S2D). These results indicate that IκBα KD preferentially affects protein phosphorylation after TNFα stimulation and attenuates the response.

To assess the temporal effects of IκBα KD on TNFα-induced protein phosphorylation, we mapped 1,286 quantified phosphosite features to kinase–substrate relationships in the human KINASE-PSP collection of PTMsigDB^40^ and used kinase–substrate enrichment analysis (KSEA)^41^ to estimate substrate-based activity scores for 80 kinases with at least five mapped phosphosites, separately for the early (0–3 h) and late (12–48 h) responses. Across the 80 kinases, baseline- referenced KSEA scores in control and IκBα KD cells were strongly correlated during the late response (Pearson *r* = 0.87), but less so during the early response (*r* = 0.42; Fig. 2D, E). Although senescence-associated phenotypes emerge after 12 h, this suggests, unexpectedly, that IκBα status has a greater effect on the early response than on the late response. In the late response, the most prominent decreases were in the inferred activities of CDK1, CDK2, CDK4, and CDK6, and these decreases were common to both control and IκBα KD cells. In contrast, in the early response, kinases involved in acute stress adaptation and survival, including AMPKA2 (an energy sensor), AKT1 (a pro-survival kinase), and MAPKAPK2 (a stress response kinase) ^42–44^, were inferred to be more strongly activated in control cells than in IκBα KD cells. These results suggest that IκBα KD does not broadly suppress TNFα-induced signaling, but preferentially blunts early adaptive stress responses.

We next examined whether the weaker early stress signaling in IκBα KD could be explained by baseline changes present before stimulation. KSEA of unstimulated cells showed that AMPKA2, AKT1, and MAPKAPK2 tended to be more active in IκBα KD cells than in control cells (Fig. 2F). Among these, AMPKA2, which is activated by reduced intracellular energy and contributes to energy homeostasis^42,45^, showed significantly higher inferred activity in IκBα KD cells (KSEA score = 3.1818, FDR = 0.016) (Fig. 2F). These results suggest that IκBα KD alters the kinase activity state before stimulation, including energy stress-related signaling, and that this elevated baseline may limit further activation after TNFα stimulation, thereby attenuating the early signaling response.

### Metabolic remodeling and mitochondrial dysfunction accompany IκBα reduction

The elevated baseline AMPKA2 activity in IκBα KD cells suggests a change in cellular energy status. Because cellular senescence is also accompanied by broad metabolic alterations, including changes in energy metabolism^46,47^, we performed metabolomic analysis of control and IκBα KD cells with or without TNFα stimulation for 48 h (Fig. 3).

**Figure 3.**
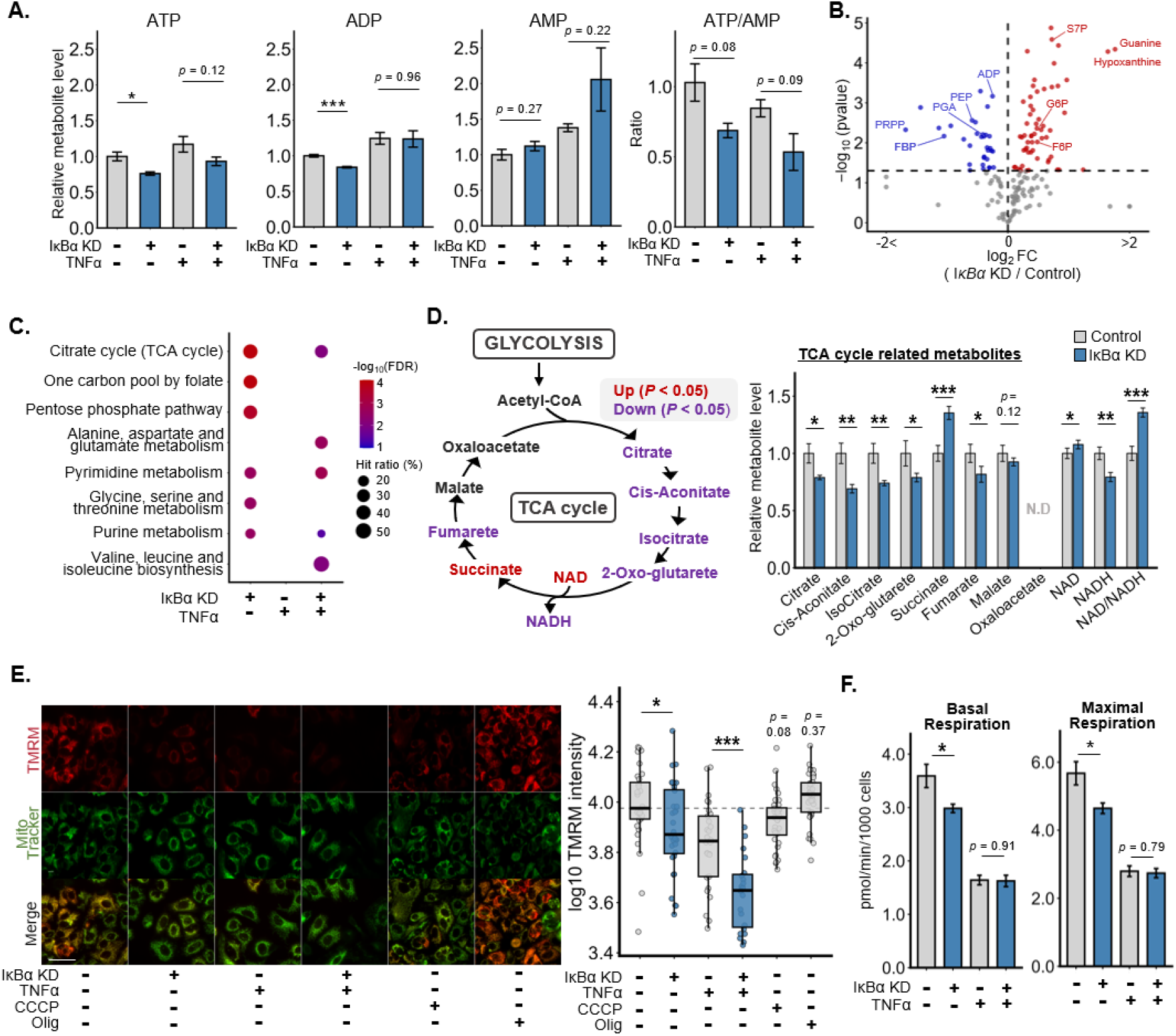
IκBα depletion remodels metabolism and impairs mitochondrial function (A) ATP, ADP, and AMP levels and the ATP/AMP ratio in Control and IκBα KD cells with or without TNFα treatment (10 ng/mL, 48 h). (B) Volcano plot of metabolite changes between Control and IκBα KD cells. Horizontal dashed line, P = 0.05. Log₂ fold changes exceeding ±2 were capped at ±2 for visualization. (C) KEGG pathway over-representation analysis of significantly altered metabolites in Control and IκBα KD cells with or without TNFα treatment (10 ng/mL, 48 h), using a hypergeometric test. Pathways containing >5 metabolites and meeting FDR < 0.05 are shown. (D) TCA cycle-related metabolite changes between Control and IκBα KD cells. Left: TCA cycle schematic showing significantly increased (red) and decreased (purple) metabolites (P < 0.05). Right: Relative metabolite levels. (E) Mitochondrial membrane potential in Control and IκBα KD cells with or without TNFα treatment (10 ng/mL, 72 h), measured by TMRM staining. Left: Representative TMRM and MitoTracker Deep Red images. Scale bars, 50 μm. Right: TMRM fluorescence intensity. Dashed line indicates the median of untreated Control cells. (F) Basal respiration (left) and proton leak (right) in Control and IκBα KD MCF-7 cells with or without TNFα treatment (10 ng/mL, 48 h), measured by Seahorse OCR analysis and normalized to cell number. For (A)–(D), metabolite levels were normalized to the mean of untreated Control cells, which was set to 1 for each metabolite. For (A)–(D), n = 4 biological replicates per group; for (E), n = 32 fields of view per group; and for (F), n = 5 per group. Bars in (A), (D), and (F) represent mean ± SEM. In (E), each point represents one field of view, and box plots show the median and interquartile range. P values in (A), (B), and (D)–(F) were determined using unpaired two-tailed Welch’s t tests for the indicated comparisons. *P < 0.05, **P < 0.01, ***P < 0.001.

In total, 168 metabolites were detected. Under unstimulated conditions, ATP and ADP levels were significantly lower in IκBα KD cells than in control cells (ATP, −23.9%, P = 0.024; ADP, −16.1%, P < 0.001) (Fig. 3A). Because AMPK is activated by a decrease in the ATP/AMP ratio^45,48^, we also calculated the ATP/AMP ratio as an indicator of cellular energy status. This ATP/AMP ratio tended to be lower in IκBα KD cells under both unstimulated (control, 1.03; IκBα KD, 0.69; P = 0.076) and TNFα-stimulated conditions (control, 0.85; IκBα KD, 0.53; P = 0.093). These results suggest that IκBα KD is associated with a lower cellular energy status, consistent with the elevated baseline AMPKA2 activity.

We next examined metabolic changes across all identified metabolites (Fig. 3B). IκBα KD increased the upstream glycolytic metabolites glucose-6-phosphate (G6P) and fructose-6-phosphate (F6P), as well as the pentose phosphate pathway intermediate sedoheptulose-7-phosphate (S7P), whereas the downstream glycolytic metabolites fructose-1,6-bisphosphate (FBP), phosphoglycerate (PGA), and phosphoenolpyruvate (PEP) decreased (P < 0.01). These changes suggest remodeling of glucose metabolism. KEGG pathway enrichment analysis of significantly altered metabolites (P < 0.05) showed that TCA cycle-related metabolites were enriched under IκBα KD alone and IκBα KD with TNFα stimulation, but not under TNFα stimulation alone (FDR = 1.7 × 10⁻⁴, 0.010, and 0.16, respectively) (Fig. 3C). Citrate, cis-aconitate, isocitrate, 2-oxoglutarate, and fumarate decreased in IκBα KD cells (P < 0.05), whereas succinate increased (log₂FC = 0.44, P < 0.001) (Fig. 3D). NAD^+^ increased and NADH decreased (P < 0.05), resulting in an increased NAD^+^/NADH ratio (log₂FC = 0.44, P < 0.001), indicating an altered intracellular redox state.

To directly assess mitochondrial function, we analyzed mitochondrial membrane potential by co-staining with TMRM and MitoTracker (Fig. 3E). IκBα KD significantly reduced TMRM intensity compared with control cells (unstimulated: −21.4%, P = 0.034; TNFα-stimulated: −36.4%, P < 0.001). The protonophore carbonyl cyanide m-chlorophenyl hydrazone (CCCP, 20 nM) and the ATP synthase inhibitor oligomycin (5 µM) were included as depolarization and hyperpolarization controls, and changed TMRM intensity by −8.2% (P = 0.075) and +13.7% (P = 0.37), respectively(Fig. 3E). We also measured mitochondrial oxygen consumption rate (OCR) using a Seahorse XF analyzer (S7801B). Although TNFα stimulation generally reduced OCR parameters, IκBα KD alone markedly impaired mitochondrial respiration even in the absence of TNFα stimulation (Fig. S3A). Specifically, IκBα KD significantly reduced basal (P = 0.032) and maximal respiration (P = 0.024) (Fig. 3F). Proton leak (P = 0.0028) and spare respiratory capacity (P = 0.032) were also significantly reduced, while ATP production showed a decreasing trend (P = 0.08) (Fig. S3B). These results indicate that IκBα KD reduces mitochondrial membrane potential and respiration, consistent with the lower ATP and ADP levels, and suggest that IκBα contributes to the maintenance of mitochondrial homeostasis.

### Mitochondrial inner and outer membrane-associated proteins are reduced in IκBα KD cells

To investigate the molecular changes underlying the mitochondrial dysfunction caused by IκBα KD, we performed time-series proteomic analysis of control and IκBα KD cells after TNFα stimulation, quantifying 8,588 proteins (see Methods). We defined organelle-associated protein sets using Gene Ontology cellular component (GOCC) annotations and performed gene set enrichment analysis (GSEA) to compare control and IκBα KD conditions at each time point. This yielded 739 mitochondrial proteins that were present in both the GOCC annotation and the quantified proteome (Fig. 4A). Among the organelle-associated protein groups examined, the mitochondrial protein group was negatively enriched in IκBα KD cells under unstimulated conditions (NES = -2.05, adjusted P = 1.1 × 10^⁻^¹⁷). Time-course analysis further showed that many of the mitochondrial proteins that were reduced by IκBα KD and remained low after TNFα stimulation were associated with the mitochondrial inner membrane. These included, under unstimulated conditions (0 h), ATPAF1 (an ATP synthase mitochondrial F1 complex assembly factor 1; log₂FC = −1.18, P < 0.001) ^49,50^, NDUFA3 (complex I subunit; log₂FC = −0.45, P = 0.0011) and NDUFB2 (complex I subunit; log₂FC = −0.62, P < 0.001)^51^, SLC25A26 (a mitochondrial S-adenosylmethionine (SAM) carrier; log₂FC = −0.41, P = 0.095)^52^, SLC25A39 (a mitochondrial glutathione transporter; log₂FC = −0.52, P < 0.001)^53^ and TIMM17A (a component of the TIM23 protein-import complex; log₂FC = −0.43, P < 0.001)^54^ (Fig. 4B). These reductions in complex I subunits and ATP synthase assembly factor may contribute to the lower ATP levels and membrane potential observed in IκBα KD cells.

**Figure 4.**
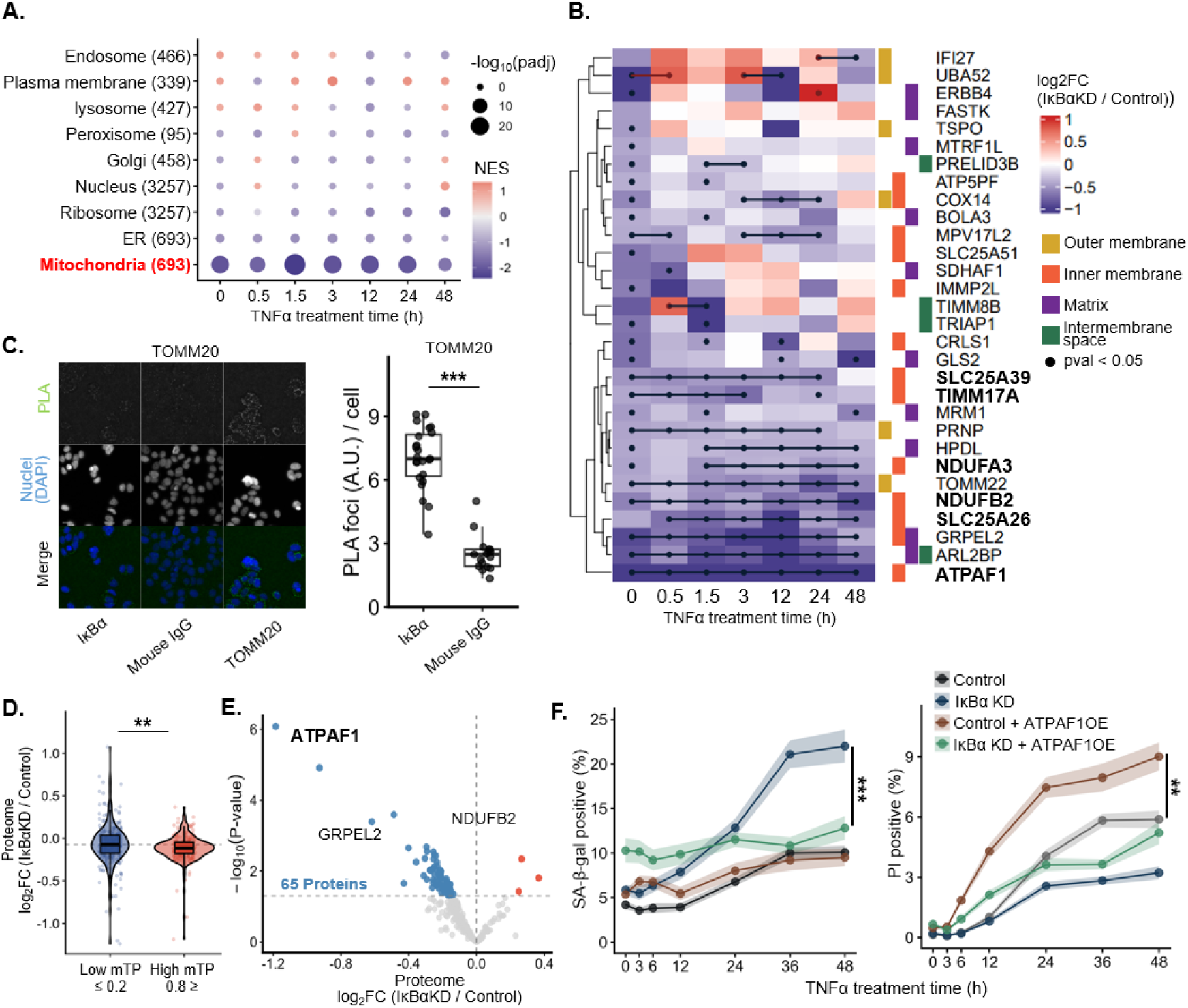
IκBα depletion links mitochondrial protein loss to senescence susceptibility (A) GSEA of GO cellular component (GOCC)-defined organelle-associated protein sets in MCF-7 cells transfected with control or IκBα siRNA (25 nM) and treated with TNFα (10 ng/mL). Proteins were ranked by logFC (IκBα KD/Control) at each time point. Numbers in parentheses indicate the number of genes included in each protein set. Dot color indicates NES, and dot size indicates −log₁₀(adjusted P value). (B) Heatmap of logFC (IκBα KD/Control) for mitochondrial leading-edge proteins. Dots indicate time points with P < 0.05 and |logFC| ≥ 0.3. Right annotations indicate mitochondrial localization based on GOCC. (C) Proximity ligation assay (PLA) of IκBα and TOMM20 in MCF-7 cells. Left: Representative PLA images. Right: Quantification of PLA foci per cell. Each point represents the mean value per image (well × field). (D) Comparison of log2FC (IκBα KD/Control) at 0 h between mitochondrial proteins with low (≤0.2) and high (≥0.8) mitochondrial targeting peptide (mTP) scores. Each point represents one protein. (E) Volcano plot of proteomic changes at 0 h (before TNFα treatment) in MCF-7 cells for mitochondrial proteins with high mTP scores (≥0.8). Proteins with P < 0.05 are shown in red (increased) or blue (decreased). (F) Time-course analysis of SA-β-gal-positive cells (left) and PI-positive cells (right) in Control and IκBα KD MCF-7 cells with or without ATPAF1 overexpression following TNFα treatment (10 ng/mL). Points indicate means, and shaded areas indicate SE. Box plots in (C) and (D) show the median and interquartile range, with whiskers extending to 1.5 × the interquartile range. P values in (C) and (F) were determined using an unpaired two-tailed Welch’s t test, and in (D) using a two-sided Wilcoxon rank-sum test. *P < 0.05, **P < 0.01, ***P < 0.001.

To ask whether the reduced mitochondrial proteins reflected lower mRNA levels, we compared mRNA and protein log₂ fold changes for mitochondrial-associated genes. Both showed a shift toward negative values in IκBα KD cells (Fig. S4A). We then classified these genes into three groups: reduced mRNA, reduced protein only, or neither (Fig. S4B), and compared ATAC-seq signals near the genes in each group (Fig. S4C). Genes with reduced mRNA tended to have lower ATAC-seq signals than genes with no reduction. Motif analysis of regions near transcription start sites (TSSs) showed enrichment of PPARA, CREB1, and YY1 motifs in this group. PPARα is a key regulator of mitochondrial energy metabolism, particularly fatty acid oxidation^55,56^. In contrast, no motifs were significantly enriched in genes with reduced protein but not mRNA (Fig. S4D), and canonical NF-κB motifs (RELA, NFKB1) were not enriched in either group. These results suggest that the reduction in mitochondrial proteins involves both transcription-associated changes and mechanisms acting after transcription.

We next explored whether mitochondrial protein import might contribute to these changes. Notably, TOMM22, a receptor subunit of the mitochondrial outer membrane translocase (TOM) complex^57^, was downregulated in unstimulated IκBα KD cells (0 h, log₂FC = −0.41, P < 0.001). Most mitochondrial proteins are encoded in the nuclear genome, synthesized in the cytosol, and imported into mitochondria through targeting signals^58^. Many of them carry a cleavable N-terminal targeting sequence (mitochondrial targeting peptide, mTP) that is recognized by TOM receptors and directs import through the TOM (outer membrane) and TIM23 (inner membrane) supercomplexes^59,60^. Because IκBα has been reported to localize to mitochondria^28,30^, we hypothesized that IκBα may support protein import at the outer membrane. To test whether IκBα is located near the TOM complex, we performed a proximity ligation assay (PLA) between IκBα and TOM subunits TOMM20, TOMM22, and TOMM40^61^ in control cells. PLA foci were most abundant with TOMM20 (7.0 vs. 2.5 foci/cell for control IgG, P < 0.001) (Fig. 4C, S5A), followed by TOMM22 (1.45 vs. 0.58 foci/cell, P < 0.001) and TOMM40 (0.29 vs. 0.12 foci/cell, P < 0.001) (Fig. S5B and S5C). TOMM20 and TOMM22 act as receptors that recognize preproteins on the cytosolic side, whereas TOMM40 forms the import channel^62,63^. These results raise the possibility that IκBα contributes to TOM complex-dependent mitochondrial protein import.

If IκBα reduction impairs import, proteins with presequences should be preferentially reduced. We therefore calculated mTP scores for each of the 739 mitochondrial proteins using TargetP^64^ and compared their abundance changes under IκBα KD between the high-(266 proteins) and low-mTP-score (290 proteins) groups (Fig. S5D). Proteins with high mTP scores decreased more than those with low scores (median log₂FC, high mTP = -0.11 vs. low mTP = -0.07; P = 0.0015) (Fig. 4D). Among the high-mTP-score proteins, 65 proteins, including ATPAF1, GRPEL2 and NDUFB2, were markedly reduced in abundance under IκBα KD compared with control cells (log₂FC < 0, P < 0.05) (Fig. 4E). These results are consistent with a regulation of IκBα to mitochondrial protein import.

We next examined whether the reduction of mitochondrial inner membrane-associated proteins in IκBα KD contributes to senescence and cell death after TNFα stimulation. For this, we overexpressed ATPAF1, an ATP synthase assembly factor and the most strongly decreased protein among the high-mTP proteins under IκBα KD (Fig. 4B), and measured the fractions of SA-β-gal-positive and PI-positive cells after TNFα stimulation (Fig. 4F). In IκBα KD cells, ATPAF1 overexpression partially suppressed the increase in SA-β-gal-positive cells (control, 10.1%; IκBα KD, 22.0%; IκBα KD + ATPAF1 OE, 12.8%; P = 0.0025 for IκBα KD vs. IκBα KD + ATPAF1 OE) and increased the fraction of PI-positive cells toward the control levels (control, 5.9%; IκBα KD, 3.2%; IκBα KD + ATPAF1 OE, 5.2%; P = 0.0039 for IκBα KD vs. IκBα KD + ATPAF1 OE). These results suggest that the reduction of ATPAF1 contributes to the senescence-prone, death-resistant phenotype of IκBα KD cells, and that impaired mitochondrial function may shift the balance between cellular senescence and cell death.

### IκBα reduction limits transition toward a pro-apoptotic state through altered mitochondrial genes

Our proteomic analysis raised the possibility that the loss of mitochondrial membrane-associated proteins accompanies the senescence-prone and death-resistant states of IκBα KD cells. However, these fractions are population-level measurements and do not show how mitochondrial state relates to senescence- or death-associated states in individual cells. To address this, we performed single-cell RNA sequencing (scRNA-seq) of control and IκBα KD cells before stimulation and after TNFα stimulation (at 6, 12, and 48 h). We then examined how mitochondrial gene expression associated with IκBα KD relates to transcriptional signatures of cellular senescence and cell death in each cell. After cell and gene filtering and downsampling to equalize cell numbers across samples (see Methods), a total of 26,608 cells were retained for downstream single-cell RNA-seq analysis. To isolate stimulus-associated transcriptional changes, we regressed out S- and G2/M-phase cell cycle scores using Seurat^65^. We then performed principal component analysis on the top 2,000 variable genes followed by graph- based clustering, and the resulting clusters were classified into six cell populations (C0–C5) for downstream analysis (Fig. 5A). Cluster composition primarily reflected time after TNFα stimulation: C0 was enriched for pre-stimulation cells, C1 and C2 for 6–12 h cells, and C3, C4, and C5 for 48 h cells (Fig. 5B). RNA velocity^66,67^ and partition-based graph abstraction (PAGA) analyses^68^ inferred a bifurcating trajectory from C0. The two branches were defined as Branch 1 (C0 → C1 → C3) and Branch 2 (C0 → C2 → C4 → C5) (Fig. 5C). Because RNA velocity infers cell-state transitions from spliced and unspliced RNA counts, we also analyzed the same data by optimal transport (OT) using the sampled time points (see Methods). This time-resolved analysis also confirmed early branching of C0 to C1 and C2 (t = 0 h to 6 h) and later transitions from C1 to C3 and C2 to C4 and C5 (t = 12 h to t = 48 h). It also identifies additional smaller contributions of transition between those clusters, that can differ between control and IκBα KD, suggesting that IκBα knockdown alters transitions toward specific cell populations (Fig. 5D). In particular, OT revealed decreased transitions from C1 to C5 after 12 h in IκBα KD condition compared to control (control, 6.3%; IκBα KD, 1.9%), which in part may explain the difference in abundance of C5 at 48 h between the two conditions. As the terminal clusters of each branch likely represent distinct stable cell fate outcomes, we compared the transcriptional features of the terminal clusters C3 and C5. Enrichment analysis of differentially expressed genes (DEGs) between these clusters showed that genes highly expressed in C3 were enriched for cell cycle-and cellular senescence-related pathways. In contrast, genes highly expressed in C5 were enriched for cell death-related pathways, including apoptosis and ferroptosis, as well as the TP53 signaling pathway (Fig. 5E). At the individual gene level, C5 showed high expression of CDKN1A, a cell-cycle arrest-associated gene, as well as stress response- and cell death-related genes, including BCL2L11, DDIT3, and GADD45A^69–72^ (Fig. 5F, S6A). Although p21 protein expression was not clearly detected by immunostaining (Supplementary Fig. 1), its mRNA was, interestingly, upregulated in cells belonging to C5, the cell death-enriched cluster.

**Figure 5.**
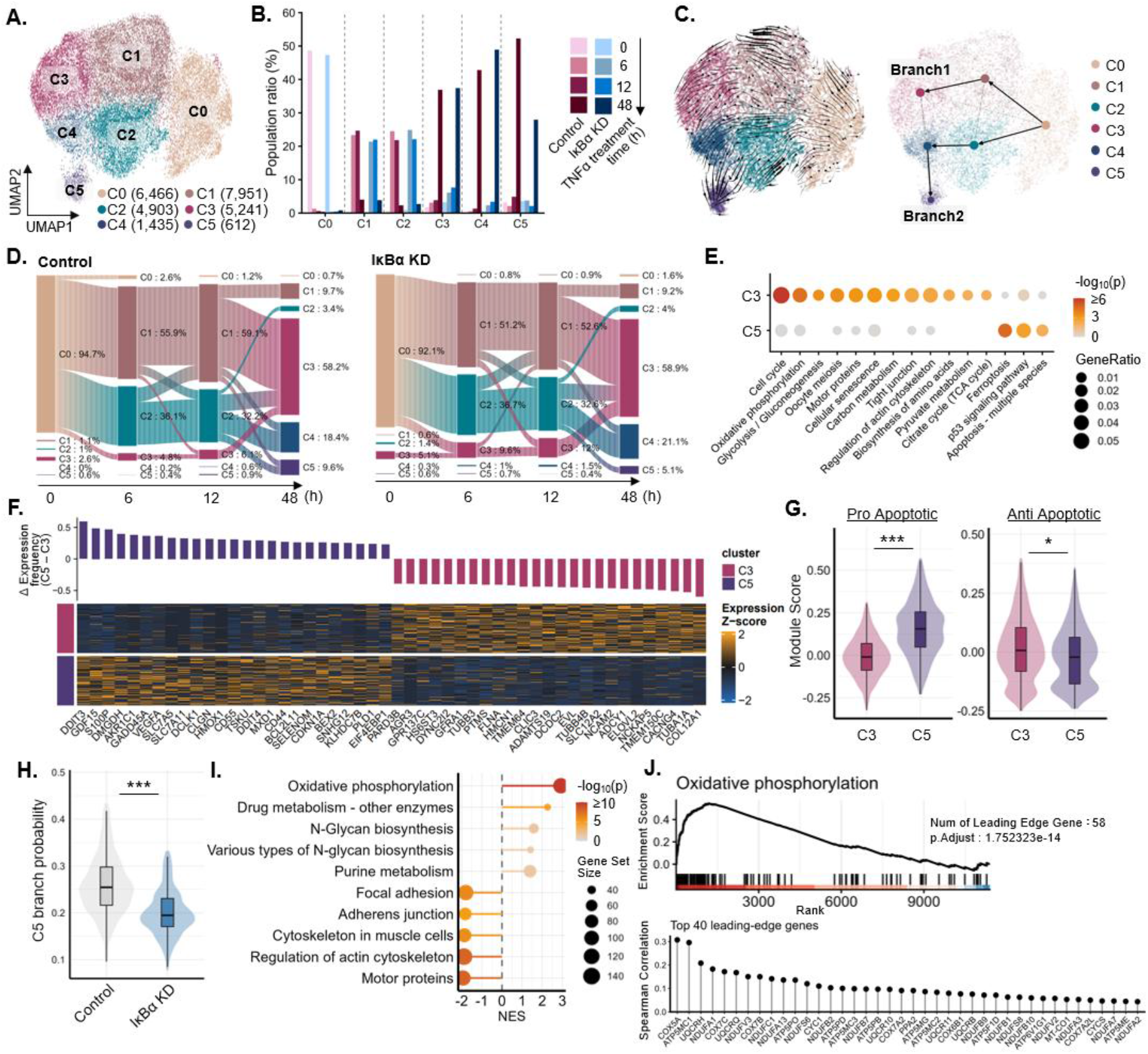
IκBα depletion lowers the predicted probability of pro-apoptotic fate commitment after TNFα stimulation (A) UMAP embedding and clustering generated using Seurat. (B) Sample-label composition of each cluster. (C) RNA velocity and PAGA analysis of cell-state transitions. Left: RNA velocity projected onto the UMAP embedding. Arrows indicate the estimated rate and direction of transcriptomic change. Right: Cluster connectivity inferred by PAGA. The trajectory from C0 through C1 to C3 was defined as Branch 1, whereas that from C0 through C2 and C4 to C5 was defined as Branch 2. (D) Sankey plots of cell-state transitions between 0–6 h, 6–12 h and 12– 48 h inferred from Optimal Transport. The bars are scaled according to cell-state abundance at each time-point. Transitions with less than 2% contribution are not shown. (E) Dot plot of KEGG enrichment analysis for differentially expressed genes between C3 and C5. (F) Heatmap of differentially expressed genes between C3 and C5. One hundred cells were randomly selected from each cluster, and Z-scored expression levels of the top 25 DEGs are shown. The top bar plot indicates the difference in the percentage of expressing cells between C5 and C3 for each gene. (G) Violin plots of module scores for pro-apoptotic BCL2 family genes (left) and anti-apoptotic BCL2 family genes (right) in C3 and C5. (H) Violin plot of C5 fate probability in C0 cells. Outliers were excluded from visualization using the 1.5 × IQR criterion. (I) Dot plot of KEGG GSEA using genes associated with C5 fate probability in C0 cells. Genes were ranked by Spearman correlation coefficients between gene expression and C5 fate probability. The five pathways with the largest absolute NES among Cellular Processes-and Metabolism-related KEGG pathways with P < 0.05 are shown. (J) GSEA of oxidative phosphorylation using genes associated with C5 fate probability. Upper: GSEA plot of oxidative phosphorylation. Lower: Lollipop plot of Spearman correlation coefficients between C5 fate probability and gene expression for the top 40 leading-edge genes. Box plots in (G) and (H) show the median and interquartile range. P values were determined using two-sided Wilcoxon rank-sum tests on 50 randomly sampled cells per group before outlier exclusion.

Since C3 and C5 diverged in cell death-related pathway enrichment (Fig. 5E), and the BCL2 family constitutes the core molecular switch governing the intrinsic mitochondrial apoptotic pathway through the balance of pro- and anti-apoptotic members^73,74^, we reasoned that the relative expression of these opposing gene groups would reflect each cluster’s susceptibility to apoptosis. To further define the C5 transcriptional state, we classified BCL2 family genes into pro-apoptotic and anti-apoptotic groups according to their established roles in apoptotic regulation ^70,75,76^, and calculated a gene score for each group. C5 showed a higher pro-apoptotic score than C3 (Median C5: 0.155; C3: −0.009; P < 0.001), whereas C3 showed a higher anti-apoptotic score than C5 (Median C3: 0.009; C5: −0.021; P = 0.020) (Fig. 5G). We next designated C3 and C5 as terminal states and used RNA velocity and CellRank^77,78^ to calculate the probability of each cell transitioning to either state by integrating RNA velocity with experimental time information (see Methods). Within the pre-stimulation C0 cluster, control-derived cells showed a higher transition probability to C5 than IκBα KD-derived cells (median estimated probability, 25.4% vs. 19.5%; P < 0.001) (Fig. 5H). These results suggest that IκBα KD reduces the likelihood of transitioning toward a pro-apoptotic transcriptional state prior to TNFα stimulation. GSEA of genes correlated with C5 branching probability revealed enrichment of oxidative phosphorylation-related pathways (NES = 2.86, adjusted P value < 0.0001) (Fig. 5I). The leading-edge genes included mitochondrial inner membrane genes involved in the electron transport chain and oxidative phosphorylation, such as COX5A^79^, ATP5MC1^80^, and UQCRH^81^ (Fig. 5J). Together, these results suggest that reduced mitochondrial gene expression associated with IκBα loss contributes to altered transcriptional-state branching following TNFα stimulation, shifting the balance between cellular senescence and cell death.

### IκB isoforms differentially affect mitochondrial homeostasis and senescence-associated phenotypes

Having established that IκBα reduction drives mitochondrial protein downregulation, SASP induction, and a shift toward the senescence-associated state, we next asked whether this phenotype is a general consequence of disrupting canonical NF-κB negative feedback, or whether it reflects a function specific to IκBα. To determine whether senescence induction associated with IκBα reduction is specific to IκBα, we individually knocked down IκBα, IκBβ, and IκBε, members of the IκB family that regulate canonical NF-κB activity^82^, and compared their phenotypes. Knockdown of each IκB isoform was confirmed by western blotting (Fig. S7A).

We first evaluated the effect of individual IκB isoform knockdown on cell morphology. Under unstimulated conditions, cell-size analysis showed that IκBα and IκBε KD induced cellular enlargement compared with control cells (both P < 0.0001), whereas IκBβ KD produced no obvious change (P = 0.978). A similar trend was observed following TNFα stimulation, with significant cellular enlargement in IκBα and IκBε KD cells (P = 0.005 and P < 0.0001, respectively), but not in IκBβ KD cells (P = 0.13). TNFα stimulation further enhanced cellular enlargement in IκBα and IκBε KD cells (both P < 0.0001) (Fig. 6A). Consistent with these morphological changes, SA-β-gal staining revealed a significant increase in the proportion of SA-β-gal-positive cells following TNFα stimulation in IκBα and IκBε KD cells (P = 0.02 and P < 0.0001, respectively) (Fig. 6B).

**Figure 6.**
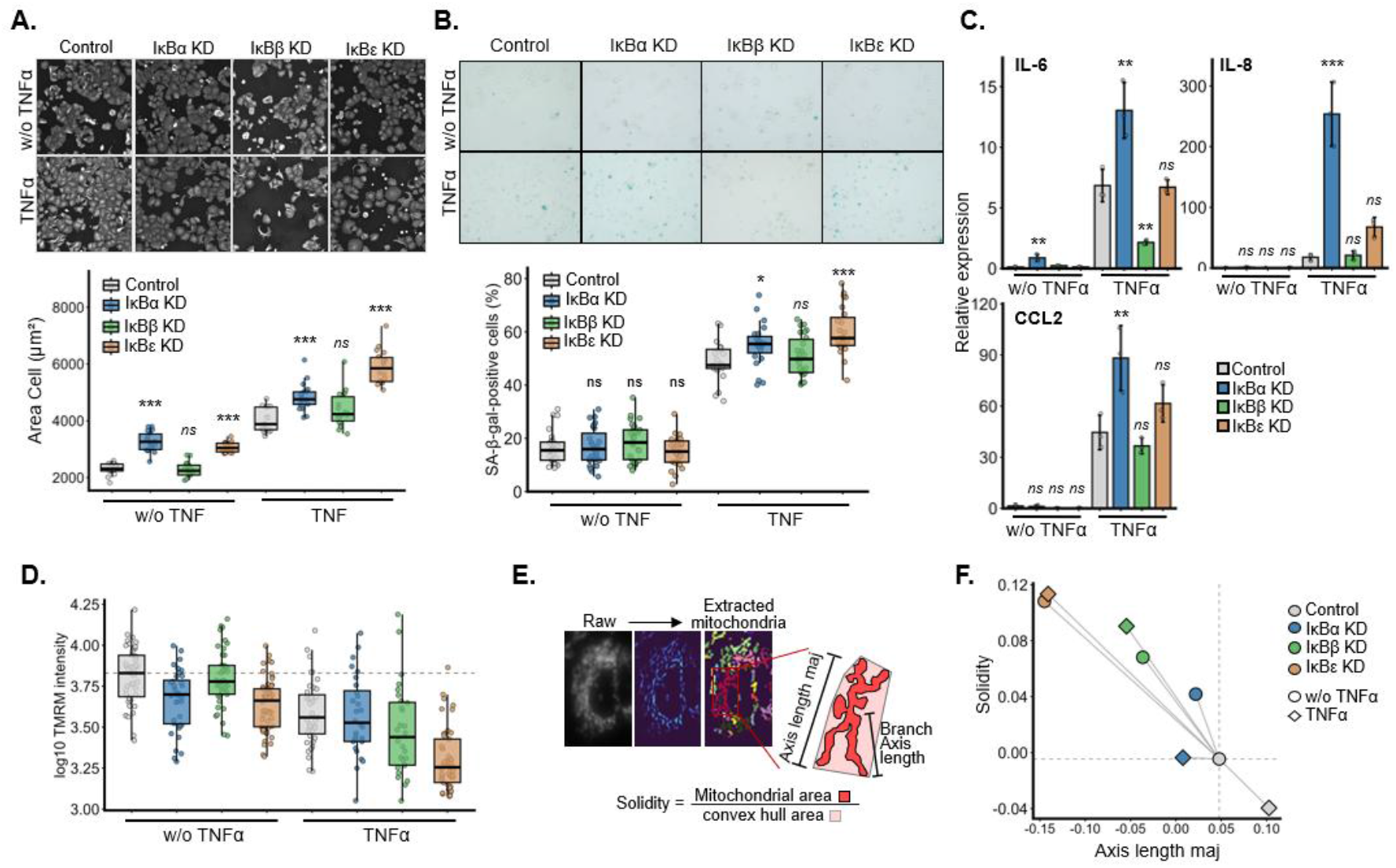
IκB isoforms differentially regulate mitochondrial homeostasis and senescence-associated phenotypes (A) Effect of IκBα, IκBβ, and IκBε KD on cell area in MCF-7 cells. Cells transfected with 25 nM Control, IκBα, IκBβ, or IκBε siRNA were treated 48 h later with TNFα (10 ng/mL), where indicated. Each point represents the mean cell area per image. (B) SA-β-gal positivity after treatment with TNFα (10 ng/mL). Representative images and quantification are shown. (C) qRT-PCR analysis of IL-6, IL-8, and CCL2 expression in MCF-7 cells transfected with 25 nM Control, IκBα, IκBβ, or IκBε siRNA and treated 48 h later with TNFα (10 ng/mL) for 48 h, where indicated. Relative expression was calculated using the 2^-ΔΔCq method with RPL27 as the internal reference. Technical replicates were averaged within each biological replicate. Bars represent mean ± SD, and each point represents one biological replicate. (D) Assessment of mitochondrial membrane potential in MCF-7 cells transfected with 25 nM Control, IκBα, IκBβ, or IκBε siRNA and treated 48 h later with TNFα (10 ng/mL) for 72 h, where indicated. Left: Representative images of TMRM and MitoTracker Deep Red fluorescence. Right: Quantification of TMRM fluorescence intensity. Each point represents one image (well × field). The dashed line indicates the median of the untreated Control condition. (E) Schematic illustration of mitochondrial morphology parameters quantified from MitoTracker images. (F) Two-dimensional comparison of mitochondrial major-axis length and solidity in MCF-7 cells transfected with 25 nM Control, IκBα, IκBβ, or IκBε siRNA and treated 48 h later with TNFα (10 ng/mL) for 72 h, where indicated. Organelle-level measurements were converted to robust z-scores relative to untreated Control cells and summarized as the median for each field. Symbols indicate condition centroids calculated as the mean of field-level median robust z-scores. Circles and diamonds indicate untreated and TNFα-treated conditions, respectively. Dashed lines indicate the untreated Control centroid, and gray lines connect the untreated Control centroid to each condition centroid. Box plots in (A),(B) and (D) show the median and interquartile range, with whiskers extending to 1.5 × the interquartile range. P values in (A)–(D) were determined separately under untreated and TNFα-treated conditions using Dunnett’s multiple comparisons test, with each KD condition compared with the corresponding Control. *P < 0.05, **P < 0.01, ***P < 0.001, ****P < 0.0001; ns, not significant.

To further assess senescence-associated inflammatory phenotypes following TNFα stimulation, we examined expression of the representative SASP factors IL6, IL8, and CCL2^16^. IκBα KD markedly increased expression of all factors (IL6, 49.88 vs. 26.1; IL8, 154.66 vs. 10.88; CCL2, 132.9 vs. 67.1; all P < 0.01), whereas IκBε KD did not clearly induce their expression, and IκBβ instead reduced IL6 expression (TNFα: 8.22 vs. 26.1, P = 0.0079) (Fig. 6C).

We next assessed mitochondrial function by measuring mitochondrial membrane potential with TMRM. Under unstimulated conditions, TMRM signals were lower in IκBα and IκBε KD cells than in control cells (3.70 and 3.66 vs. 3.83, respectively; both P < 0.001) (Fig. 6D). Analysis of mitochondrial morphology further revealed that mitochondrial shortening and fragmentation were stronger under IκBβ and IκBε KD than under IκBα KD (Fig. 6E,F, S7B-D). Notably, Western blot analysis revealed a reduction in ATPAF1 protein levels specifically in IκBα KD cells, but not in IκBβ or IκBε KD cells (P < 0.0001, P = 0.32 and 0.99, respectively) (Fig. S8A).

Taken together, these results revealed a clear dissociation among IκB isoforms: IκBα KD produced the most robust senescence-associated phenotype (cellular enlargement, SA-β-gal and SASP induction) despite causing the mildest mitochondrial disruption whereas IκBβ and IκBε caused more severe mitochondrial dysfunction without a corresponding increase in SASP expression of cellular enlargement. This indicates that IκB isoforms are not functionally interchangeable within the canonical NF-κB pathway, and the extent of mitochondrial dysfunction alone does not predict whether a senescence-associated inflammatory phenotype is induced. These findings suggest that, beyond their shared role in NF-κB suppression, IκB isoforms have distinct, non-overlapping functions linking mitochondrial regulation to downstream cellular outcomes, with IκBα uniquely coupling its loss to a senescence-associated transcriptional program.

## DISCUSSION

NF-κB signaling is a central regulatory system for inflammatory responses and cell survival, and IκBα, a canonical inhibitor of NF-κB, has primarily been understood as a controller of NF-κB-dependent transcriptional responses^4,7,83^. Our findings suggest that IκBα also contributes to mitochondrial homeostasis and post-stress cell-fate decisions. IκBα reduction appears to act as a priming event that shifts cells toward a stress-responsive state prone to senescence, rather than directly inducing a senescent state. Even without TNFα stimulation, IκBα reduction caused impaired mitochondrial homeostasis, including lower ATP levels, reduced respiratory function, and reduced levels of mitochondrial proteins, including those involved in protein import.

Because mitochondria regulate not only ATP production but also cell death and stress adaptation^84–86^, these changes may influence how cells respond to subsequent damage. The resulting lower energy level may also relate to the weaker kinase responses to TNFα, which could further affect fate decisions controlled by the signaling network.

Senescence and cell death are both protective responses to damaged cells, but senescent cells often maintain survival while exhibiting inflammatory phenotypes^19,87^, and they depend on anti- apoptotic pathways^39,88^. In IκBα KD cells, post-stimulation cell death was suppressed and branching toward the pro-apoptotic state was reduced, consistent with a survival-biased fate. These findings suggest that mitochondrial changes accompanying IκBα reduction may bias cell- fate selection away from death and toward senescence. IκBα reduction has been reported in chronic inflammatory environments and age-associated cellular states^21,89^, this relationship may help explain the accumulation of senescent cells and altered inflammatory responses during aging.

How IκBα contributes to mitochondrial homeostasis remains an open question. IκBα and NF-κB have been suggested to localize to mitochondria^28,29^, and a recent study reported that forced mitochondrial targeting of IκBα in lung cancer cells promotes metabolic rewiring, and metastasis^90^. Our observation that IκBα localizes near the TOM complex and appears to support maintenance of proteins bearing mitochondrial targeting signals suggests that IκBα reduction may directly impair mitochondrial function through a noncanonical, import-associated mechanism. This function appears distinct from, or to operate in parallel with, the canonical role of IκBα in NF-κB-dependent transcriptional regulation. Knockdown of IκBβ and IκBε also affected mitochondrial morphology, suggesting that this role may extend to other IκB family members, although their effects on senescence-associated phenotypes differed from those of IκBα. Future studies should test this possibility using protein-level validation at single-cell resolution, super-resolution imaging, and rescue experiments that manipulate IκB localization and mitochondrial morphologies.

Overall, these findings position IκBα not merely as a negative regulator of NF-κB transcription, but as a node linking mitochondrial homeostasis to post-stress cell fate decisions. This framework offers a mechanistic link between chronic inflammation, metabolic decline, and senescent-cell accumulation, and suggests that IκB family members more broadly warrant attention as determinants of cell fate during aging and chronic diseases.

## RESOURCE AVAILABILITY

### Lead contact

Requests for raw data and code should be directed to, and will be fulfilled by the Lead Contact, Mariko Okada.

## Materials availability

This study did not generate new unique reagents.

## Data and code availability

All omics data will be available in public repositories upon publication. The code for the bioinformatics analysis and mathematical modeling will also be available on GitHub (https://github.com/okadalabipr/Matsuda2026) upon publication.

Any additional information required to reanalyze the data reported in this paper is available from the lead contact upon request.

## ACKNOWLEDGMENTS

We thank Dr. Josephina Sampson for critical reading of the manuscript and for supervision of some of experiments. We would also like to thank Dr. Hsueh-Fen Juan, Dr. Hsuan-Cheng Huang, Ms. Tang-Ching Tang and Mr. Tzu-Yang Tseng for their discussions on proteomics analysis and cellular senescence. This study was supported in part by JSPS Bilateral Programs (JPJSBP120245703), JST ASPIRE Program (JPMJAP24B1), JST CREST Program (JPMJCR21N3), and JST ARiSE Program (JPMJAR2623) to M.O., and JSPS KAKENHI (JP26K09267) to K.H.. This research was also partially supported by the Research Support Project for Life Science and Drug Discovery (Basis for Supporting Innovative Drug Discovery and Life Science Research [BINDS]) from AMED under Grant Numbers JP22ama121055 to T.B. and JP25ama121055 to S.T.. This work was performed in part under the International Collaborative Research Program of the Institute for Protein Research, Osaka University (ICR-26-8).

## AUTHOR CONTRIBUTIONS

S.S., Sho T., T.K. and H.H. performed cell-based experiments, and T.K. and H.H. analyzed cell morphology; K.H. and A.H. performed functional analysis of mitochondria; J.A. acquired the proteome and phosphoproteome data and performed the primary data analysis; Y.I. and T.B. acquired the metabolome data and performed the primary data analysis; Shashank T., B.G. and J.W. performed optimal transport analysis of scRNA-seq data; K.M. performed the integrative computational analysis of all omics data. K.M. and M.O. interpreted the results and wrote the original draft and the final manuscript; M.O. conceived and supervised the study. All authors reviewed and approved the final manuscript.

## DECLARATION OF INTERESTS

The authors declare that they have no competing interests.

## DECLARATION OF GENERATIVE AI AND AI-ASSISTED TECHNOLOGIES

During the preparation of this work, the authors used Anthropic Claude and OpenAI ChatGPT in order to edit and fix the sentences and search for relevant literature. After using this tool, the authors reviewed and edited the content as needed and take full responsibility for the content of the publication.

## METHOD DETAILS

### Cell culture

Human breast adenocarcinoma MCF-7 cells (American Type Culture Collection) were cultured in Dulbecco’s modified Eagle’s medium (DMEM; Nacalai Tesque) supplemented with 10% (v/v) fetal bovine serum (FBS; Sigma-Aldrich) and antibiotics (100 U/mL penicillin, 100 μg/mL streptomycin, and 0.25 μg/mL amphotericin B; Nacalai Tesque, Kyoto, Japan) at 37 °C in a humidified atmosphere containing 5% CO₂.

### siRNA transfection and cell treatment

For siRNA transfection, siRNAs targeting IκBα (ON-TARGETplus SMARTpool, L-004765-00), IκBβ (ON-TARGETplus SMARTpool, L-015757-00), IκBε (ON-TARGETplus SMARTpool, L- 004766-00), RELA (ON-TARGETplus SMARTpool, L-003533-00), and ATPAF1 (ON-TARGETplus SMARTpool, L-013738-01), together with a negative-control siRNA (D-001810- 02), were purchased from Dharmacon/Horizon Discovery (Cambridge, UK). Cells were transfected with siRNA at a final concentration of 25 nM using Lipofectamine RNAiMAX reagent (Thermo Fisher Scientific) in serum-reduced Opti-MEM (Thermo Fisher Scientific), according to the manufacturer’s instructions. For stress-treatment assays, cells were treated 48 h after siRNA transfection with TNFα (10 ng/mL; Thermo Fisher Scientific), H₂O₂ (300 μM; Wako), etoposide (10 μM; Sigma-Aldrich), or camptothecin (10 μM; Sigma-Aldrich). For TNFα washout experiments, cells were stimulated with TNFα after siRNA transfection, and TNFα-containing medium was replaced with TNFα-free medium after 0, 3, 6, 12, 24, or 48 h of stimulation. Cells were then cultured in TNFα-free medium until 48 h after the initial TNFα addition.

### Plasmid transfection for gene over-expression

The expression plasmid encoding human ATPAF1 (pRP[Exp]-EGFP/Puro- CAG>hATPAF1[NM_022745.4], vector ID: VB900017-5491hqu) was purchased from VectorBuilder (Chicago, IL, USA). Cells were seeded in 24-well plates at a density of 1 × 10⁵ cells/well 24 h before transfection. Transfection was performed using 0.1 µg of plasmid DNA and Lipofectamine LTX Reagent without PLUS Reagent (Thermo Fisher Scientific, Waltham, MA, USA) according to the manufacturer’s instructions. At 24 h after transfection, cells were subjected to TNFα treatment.

### Western blotting

Western blot analysis was performed as described previously^91^ using primary antibodies against GAPDH (Sigma-Aldrich; #MAB374; 1:2,000), IκBα (Cell Signaling Technology; #4814; 1:1,000), anti-GRPEL2 rabbit antibody (Proteintech; #17751-1-AP), anti-mtHSP70 rabbit antibody (Cell Signaling Technology; #3593S), anti-IMMP2L rabbit antibody (Proteintech; #PGI-15970-1-AP- 150), anti-ANT1 rabbit antibody (Cell Signaling Technology; #51755S), anti-ANT1 mouse antibody (Santa Cruz Biotechnology; #SCB-SC-518268-200), anti-ATPAF1 mouse antibody (Santa Cruz Biotechnology; #SC-398684), and anti-ATPAF1 rabbit antibody (Invitrogen; #PA5- 31394). HRP-linked anti-mouse IgG (Cell Signaling Technology; #7076; 1:5,000) or HRP-linked anti-rabbit IgG (Cell Signaling Technology; #7074; 1:5,000) were used as secondary antibodies.

### Quantitative reverse transcription PCR (qRT-PCR)

Total RNA was isolated from cells using the NucleoSpin RNA Plus kit (Macherey-Nagel; #U0984C) according to the manufacturer’s instructions. cDNA was synthesized from 1 μg of total RNA using ReverTra Ace qPCR RT Master Mix (TOYOBO; #FSQ-201) according to the manufacturer’s instructions. qRT-PCR was performed using KOD SYBR qPCR Mix (TOYOBO; #QKD-201X5) with a CFX96 Real-Time PCR Detection System (Bio-Rad). Each 20 μL PCR reaction contained 10 μL KOD SYBR qPCR Mix, 2 μL of forward primer (2 μM), 2 μL of reverse primer (2 μM), 2 μL of cDNA, and 4 μL of distilled water. PCR amplification was carried out for 40 cycles under the following conditions: 98 °C for 10 s, 60 °C for 10 s, and 68 °C for 30 s. The primer sequences used for qRT-PCR are listed in Table 1. Relative gene expression was calculated using the 2^−ΔΔCq^ method with RPL27 as the internal reference gene. Each independent experiment was performed in triplicate.

### Senescence-associated β-galactosidase (SA-β-gal) staining

Senescent cells were stained using a Senescence β-Galactosidase Staining Kit (Cell Signaling Technology; #9860), according to the manufacturer’s instructions and as described previously^21^. Briefly, cells in 24-well plates were washed with PBS, fixed for 15 min in fixative solution, washed twice with PBS, and incubated in SA-β-gal staining solution overnight at 37 °C. Images were acquired with a Keyence BZ-9000 microscope at 40× magnification. Four randomly selected fields per well were used to count SA-β-gal-positive cells. The percentage of SA-β-gal- positive cells was analyzed using ImageJ. DAPI staining was used to identify nuclei and count the total number of cells per field.

### Cell staining

Cell viability was assessed by double staining with Hoechst 33342 and propidium iodide (PI) according to the manufacturers’ protocols. Cells were incubated with Hoechst 33342 (Nacalai Tesque, #04929-82; 5 µg/mL) and PI (Nacalai Tesque, #19174-31; 1 µg/mL) for 10 min at room temperature, protected from light. To retain loosely attached dead cells, no washing step was performed, and cells were imaged without fixation. Images were acquired using an IN Cell Analyzer 2500HS (Cytiva, Marlborough, MA, USA).

### Elution-based iterative indirect immunofluorescence (4i) and high-content imaging

Cells were reverse-transfected with siRNAs as described above and replated into 96-well imaging plates at 12,000, 10,000, 7,000, or 4,000 cells per well for the 0-, 24-, 48-, or 72-h experimental endpoints, respectively. After overnight incubation, cells were stimulated with the indicated reagents at the specified concentrations and incubated for the indicated durations at

37 °C in a humidified atmosphere containing 5% CO₂. For EdU incorporation assays, cells were incubated with 50 μM 5-ethynyl-2-deoxyuridine (EdU) for 1 h before fixation. Cells were fixed with 4% paraformaldehyde for 20 min at room temperature and permeabilized with 0.1% Triton X-100 in PBS for 15 min. 4i staining was based on the elution protocol described by Gut et al.^34^. The protocol adaptations comprised 30-min primary and secondary antibody incubations at room temperature, use of conventional blocking solution (cBS), SYTOX Green counterstaining, and EdU click detection in the final round. Briefly, cells were incubated with primary antibodies diluted in conventional blocking solution (cBS) for 30 min at room temperature, washed with PBS, and subsequently incubated with Alexa Fluor 647-conjugated anti-mouse and Alexa Fluor 555-conjugated anti-rabbit secondary antibodies (Thermo Fisher Scientific; 1:500) for 30 min at room temperature. Nuclei were counterstained with SYTOX Green (1:10,000 dilution in TBS) for 15 min at room temperature. Images were acquired in imaging buffer (IB) using an Operetta CLS High-Content Analysis System (PerkinElmer). After each imaging cycle, fluorescence signals were removed by three consecutive 10-min incubations with elution buffer at room temperature. Complete signal removal was confirmed by re-imaging the cells after elution.

Subsequent staining rounds were performed by repeating the staining, imaging, and elution procedures. EdU incorporation was detected in the final staining round using the Click-iT™ Plus EdU Cell Proliferation Kit for Imaging, Alexa Fluor™ 647 dye (Thermo Fisher Scientific; #C10640), according to the manufacturer’s instructions. OMERO-Screen performed flatfield correction, Cellpose-based nuclear and cell segmentation, and per-cell feature extraction, including nuclear and cytoplasmic areas, nuclear DNA content, EdU incorporation. Cross-round registration used the SYTOX Green channel, followed by cell matching and alignment quality control. The workflow is documented at https://hocheggerlab.github.io/omero-screen/cyclic-if/, and the software source is available in the HocheggerLab/omero-screen GitHub repository (https://github.com/HocheggerLab/omero-screen). Primary antibodies used for 4i included antibodies against IκBα (Cell Signaling Technology; #4814; 1:500), p21 (Cell Signaling Technology; #2947; 1:500), α-tubulin (Cell Signaling Technology; #3873; 1:1,000), phospho-Rb (Ser807/811; Cell Signaling Technology; #8516; 1:500), p53 (Cell Signaling Technology; #48818; 1:500), and Cyclin A2 (Cell Signaling Technology; #67955; 1:500). Cytoplasmic enlargement was quantified at the single-cell level using control cells before stimulation as the reference. A robust linear regression model was fitted between log₁₀-transformed nuclear and cytoplasmic areas. Cells with residuals above the 99th percentile of the reference residual distribution were defined as cytoplasmically enlarged. The percentage of enlarged cells was calculated for each image and summarized as the mean ± SEM across images.

### Immunofluorescence staining of p27 and α-tubulin

Cells were fixed with 4% paraformaldehyde (PFA; Thermo Fisher Scientific; #28908) for 15 min at room temperature and washed twice with PBS(-). Cells were then permeabilized with 0.1% Triton X-100 (Nacalai Tesque; #35501-15) for 15 min at room temperature with gentle shaking, followed by two washes with PBS(-). After permeabilization, cells were blocked with 10 mg/mL bovine serum albumin (BSA; Nacalai Tesque; #01859-47) for 1 h at room temperature with gentle shaking. Primary antibodies were diluted in BSA blocking buffer and incubated with the cells overnight at 4 °C with gentle shaking. The primary antibodies used were mouse monoclonal anti-α-tubulin (Cell Signaling Technology; #3873; 1:1,000) and rabbit monoclonal anti-p27 Kip1 (D69C12) (Cell Signaling Technology; #3686; 1:500). Following primary antibody incubation, cells were washed three times with PBS(-) for 10 min each with gentle shaking.

Cells were subsequently incubated with goat anti-mouse IgG Alexa Fluor 488 (Thermo Fisher Scientific; #A11001; 1:500) and donkey anti-rabbit IgG DyLight 650 (Thermo Fisher Scientific; #SA5-10041; 1:500), diluted in BSA blocking buffer, for 1 h at room temperature with gentle shaking in the dark. Cells were then washed three times with PBS(-) for 10 min each in the dark. Nuclei were stained with 1 μg/mL DAPI (Nacalai Tesque; #11034-56) for 15 min at room temperature with gentle shaking in the dark, followed by two washes with PBS(-). Fluorescence images were acquired using an IN Cell Analyzer 2500HS (GE Healthcare).

### Cell morphology analysis by high-content imaging

Cells were seeded into 96-well imaging plates at a density of 1 × 10⁴ cells/well. After overnight incubation, cells were stimulated with TNFα (10 ng/mL) and incubated for 48 hours at 37 °C in a humidified atmosphere containing 5% CO₂. Cells were fixed with 4% paraformaldehyde for 15 min at room temperature and permeabilized with 0.1% Triton X-100 in PBS for 15 min.

Subsequently, cells were stained with CellMask™ Plasma Membrane Stains (Thermo Fisher Scientific, Cat. No. C10046; 100 ng/mL) and 4′,6-diamidino-2-phenylindole dihydrochloride (DAPI; Nacalai Tesque, Cat. No. 11034-56; 1 µg/mL) for 15 min. Images were acquired using an IN Cell Analyzer 2500HS (Cytiva, Marlborough, MA, USA).

### Analysis of mitochondrial morphology and membrane potential

MCF-7 cells were seeded at 1.5 × 10⁴ cells per well in 96-well clear-bottom plates (PerkinElmer; #6055302). Unless otherwise specified, cells were stained 18 h after seeding with 100 nM tetramethylrhodamine methyl ester (TMRM; Setareh Biotech; #SEB-6275-25), 100 nM MitoTracker Deep Red FM (Cell Signaling Technology; #8778), and 1 μg/mL Hoechst 33342 (Dojindo; #346-07951) for 30 min at 37 °C and 5% CO₂. After washing with phenol red-free DMEM (FUJIFILM; #044-32955) supplemented with 10% (v/v) FBS (NICHIREI BIOSCIENCES

INC.; #174012), cells were incubated in phenol red-free medium at 37 °C for 30 min and then imaged at 37 °C and 5% CO₂. As positive and negative controls for TMRM-based mitochondrial membrane potential measurements, cells were treated with 5 μM oligomycin and 20 nM carbonyl cyanide m-chlorophenyl hydrazone (CCCP), respectively. Imaging was initiated 5 min after oligomycin treatment and 2 h after CCCP treatment. Single-optical-section images were acquired using an IN Cell Analyzer 2500HS (GE Healthcare) equipped with a CFI Plan Fluor ELWD 40× dry objective lens (NA 0.60). MitoTracker Deep Red was imaged using excitation wavelengths of 621–643 nm and emission wavelengths of 662–700 nm; TMRM using excitation wavelengths of 528.5–555.5 nm and emission wavelengths of 574.5–619.5 nm; and Hoechst 33342 using excitation wavelengths of 381–399 nm and emission wavelengths of 411–459 nm. Mitochondrial morphology and fluorescence intensity were quantified using Nellie^92^. For morphology analysis, organelle-level measurements of mean branch length, mean branch tortuosity, major-axis length, and solidity were used. Branch length and major-axis length were log₁₀-transformed, and organelles lacking valid measurements for any of the four parameters were excluded. For each parameter, robust z-scores were calculated relative to untreated control cells using the median and median absolute deviation (MAD; scaling factor, 1.4826) and capped at ±5. Organelle-level robust z-scores were summarized as the median for each field, and fields containing fewer than 20 organelles were excluded. For two-dimensional analysis of major-axis length and solidity, condition centroids were calculated as the mean of the corresponding field-level median robust z-scores. Data processing, statistical analysis, and visualization were performed using R.

### Proximity ligation assay (PLA)

MCF-7 cells were seeded at 1.5 × 10⁴ cells per well in 96-well clear-bottom plates (PerkinElmer; #6055302). PLA was performed according to the manufacturer’s protocol using Duolink In Situ Detection Reagents Green (Sigma-Aldrich; #DUO92014-100RXN) and PLA probes (Anti-Rabbit MINUS; #DUO92005-100RXN; Anti-Mouse PLUS; #DUO92001-100RXN). Unless otherwise specified, cells were fixed 18 h after seeding with 4% formaldehyde (Thermo Fisher Scientific; #28908) for 15 min at room temperature and permeabilized with 0.5% Triton X-100 (Sigma- Aldrich; #T-9284) for 5 min. After washing three times with D-PBS (Nacalai Tesque; #14249- 24), cells were incubated with blocking solution for 1 h at 37 °C. Primary antibodies were diluted 1:500 and incubated with the cells overnight at 4 °C. Rabbit IgG (Cell Signaling Technology; #3900) and mouse IgG (Cell Signaling Technology; #5415) were used as negative controls. The following primary antibodies were used: anti-IκBα mouse antibody (Cell Signaling Technology; #4814S), anti-IκBα rabbit antibody (Abcam; #ab76429), anti-IκBβ rabbit antibody (Cell Signaling Technology; #15519), anti-IκBε rabbit antibody (Santa Cruz Biotechnology; #SC-7156), anti- TOMM20 rabbit antibody (Abcam; #ab186735-100), anti-TOMM20 mouse antibody (Santa Cruz Biotechnology; #SCB-SC-17764-200), anti-TOMM22 rabbit antibody (Cell Signaling Technology; #90704), and anti-TOMM40 mouse antibody (Santa Cruz Biotechnology; #SCB- SC-365467-200). After washing twice with Buffer A, diluted PLA probes were added and incubated for 1 h at 37 °C. Cells were then washed twice with Buffer A, and the ligation reaction was performed for 30 min at 37 °C. After two additional washes with Buffer A, rolling circle amplification was performed for 100 min at 37 °C, followed by washing with Buffer B. Finally, cells were mounted using Duolink In Situ Mounting Medium with DAPI (Sigma-Aldrich; #DUO82040-5ML). Single-optical-section images were acquired using an IN Cell Analyzer 2500HS (GE Healthcare) equipped with a CFI Plan Fluor ELWD 40× dry objective lens (NA 0.60). PLA signals were imaged using excitation wavelengths of 461–489 nm and emission wavelengths of 499.5–522.5 nm, and DAPI using excitation wavelengths of 381–399 nm and emission wavelengths of 411–459 nm. Images were processed using ImageJ version 1.54p (https://imagej.net/ij/index.html) to enhance contrast and reduce background haze. PLA foci and nuclei were then counted.

### Oxygen consumption rate (OCR) assay

Oxygen consumption rate (OCR) was measured using a Seahorse XFe24 Analyzer (Agilent Technologies, S7801B). Cells were seeded in Seahorse XFe24 Cell Culture Microplates (Agilent Technologies; FluxPak XFe24 Assay Kit; #102340-100). Before measurement, the culture medium was replaced with XF DMEM medium (Agilent Technologies; #103575-100) supplemented with 1 mM pyruvate (Agilent Technologies; #103578-100), 2 mM glutamine (Agilent Technologies; #103579-100), and 10 mM glucose (Agilent Technologies; #103577- 100). Cells were incubated for approximately 60 min at 37 °C in a non-CO₂ incubator to allow temperature and pH equilibration. The sensor cartridge was hydrated with XF Calibrant (Agilent Technologies; #100840-000) according to the manufacturer’s instructions. OCR was determined using the Seahorse XF Cell Mito Stress Test Kit (Agilent Technologies; #103015-100).

Oligomycin (1.5 μM), FCCP (1.0 μM), and rotenone/antimycin A (0.5 μM) were sequentially injected during the assay. Basal respiration, ATP production, maximal respiration, and spare respiratory capacity were calculated from the OCR profiles using Agilent Seahorse XFe Wave software. After the assay, cells were stained with Hoechst 33342, and nuclei were imaged and automatically counted using a Cytation 1 Cell Imaging Multi-Mode Reader (Agilent Technologies). OCR values were normalized to the cell number.

### Proteome sample preparation

Each sample was solubilized with phase transfer surfactant (PTS) buffer^93^. A total of 200 μg of each sample was transferred to a 1.5 mL tube and boiled at 95 °C for 5 min. Samples were reduced with 10 mM TCEP, alkylated with 20 mM iodoacetamide, and proteins were purified using the SP3 protocol ^94^. Purified proteins were digested with trypsin and Lys-C (each at an enzyme-to-protein ratio of 1:50, w/w) for 16 h at 37 °C. For proteome analysis, 2.5% of the digested peptides were desalted and purified using C18-SCX StageTips ^95^. Peptides were dried in a SpeedVac and reconstituted in 1% trifluoroacetic acid/2% acetonitrile.

The remaining digested peptides were desalted and applied to an IMAC column for phosphopeptide enrichment ^96^. TMTpro 16plex reagents (0.5 mg; Thermo Scientific) were dissolved in 40 μL of anhydrous acetonitrile, and 4 μL of this solution was added to phosphopeptides dissolved in 10 μL of 100 mM TEAB. After incubation for 1 h at room temperature, the reaction was quenched with hydroxylamine to a final concentration of 0.3% (v/v). TMT-labeled phosphopeptides were subjected to off-line basic pH reversed-phase (BPRP) fractionation^97^.

### Phosphopeptide fractionation by basic pH reversed-phase HPLC

The pooled TMT-labeled peptide sample was fractionated by BPRP HPLC using an UltiMate 3000 UHPLC system (Thermo Scientific) equipped with a dual-wavelength detector set at 220 and 280 nm. Mobile phases were BPRP-A (5 mM ammonium bicarbonate pH 9.2) and BPRP-B (5 mM ammonium bicarbonate pH 9.2 and 60% acetonitrile). The LC gradient for total proteomics samples was increased from 5% to 25% BPRP-B over 4 min, from 25% to 60% over 26 min, and from 60% to 99% over 5 min. TMT-labeled phosphopeptides were separated using an L-column3 C18 column (5 μm particles, 0.3 mm ID and 150 mm in length, Chemicals Evaluation and Research Institute, Japan) at a flow rate of 2 μL/min. The peptide mixture was fractionated into 21 fractions, which were consolidated into 7 fractions. Samples were subsequently vacuum-centrifuged to near dryness. Each fraction was reconstituted in 2% acetonitrile, 0.1% trifluoroacetic acid for LC-MS/MS analysis.

### Proteome LC-MS/MS analysis

LC-MS/MS was performed by coupling an UltiMate 3000 Nano LC system (Thermo Scientific, Bremen, Germany) and an HTC-PAL autosampler (CTC Analytics, Zwingen, Switzerland) to an Orbitrap Fusion Lumos mass spectrometer (Thermo Scientific). The Orbitrap Fusion Lumos instrument was operated in the data-independent acquisition (DIA) mode. Gas-phase fractionation (GPF)-DIA acquisition was done using a pooled sample for library, Full mass spectra were acquired with the following parameters: a 2 *m/z* isolation window, a resolution of 120,000, an automatic gain control (AGC) target 250 %with custom max injection time, and a normalized collision energy of 30. The five GPF-DIA runs collectively covered 420-780 *m/z* (i.e., 418-494, 490-566, 562-638, 634-710, and 706--782 *m/z*). MS2 spectra were collected with the following parameters: a 2-*m/z* isolation window at 50,000 resolution, an AGC target of 2 × 10^5^ ions, a maximum injection time is 86 ms, and normalized collision energy of 30. For the individual samples for proteome profiling acquisition, full mass spectra were acquired in the range of 420–780 *m/z* with the following parameters: a resolution of 120,000, an AGC target of 4 × 10^5^ with a maximum injection time of 100 ms. MS2 spectra were collected with the following parameters: a 10-*m/z* isolation window at 30,000 resolution, an AGC target of 2 × 10^5^ ions, a maximum injection time is 54 ms, overlapping window patterns and normalized collision energy of 30.

### Proteome data preprocessing

Raw MS data were processed with DIA-NN version 1.8. Database searching included all entries from the Homo sapiens UniProt (2020/04 release) database and contaminant database ^98^. The search parameters were as follows: up to two missed cleavages, peptide lengths of 7–30 amino acids, carbamidomethylation of cysteine residues (+57.021 Da) as a fixed modification, protein names from FASTA for implicit protein grouping, robust LC (high precision) for quantification, and retention time-dependent cross-run normalization. Precursor identifications were filtered at a false discovery rate (FDR) of 1%. MaxLFQ was used for label-free quantification.

Filtering and missing value imputation of the acquired data were performed using Perseus version 2.0.10.0. Proteins quantified in all three replicates at one or more time points in either condition were retained for subsequent analysis. After filtering, remaining missing values were imputed with random values drawn from a normal distribution downshifted by 1.8 standard deviations from the distribution of observed values, with a width of 0.3, according to the method described by Tyanova et al.^99^.

### Phosphoproteome LC-MS/MS analysis

LC-MS/MS was performed by coupling an UltiMate 3000 Nano LC system (Thermo Scientific, Bremen, Germany) and an HTC-PAL autosampler (CTC Analytics, Zwingen, Switzerland) to a Q Exactive plus hybrid quadrupole-Orbitrap mass spectrometer (Thermo Scientific).

Phosphopeptides were delivered to an analytical column (75 μm × 30 cm, packed in-house with ReproSil-Pur C18-AQ, 1.9 μm resin, Dr. Maisch, Ammerbuch, Germany) and separated at a flow rate of 280 nL/min using a 145-min gradient from 5% to 30% of solvent B (solvent A, 0.1% formic acid (FA); solvent B, 0.1% FA and 99.9% acetonitrile). The Orbitrap Fusion Lumos instrument was operated in the data-dependent acquisition (DDA) mode. For global phosphoproteome analysis, survey full scan MS spectra (*m/z* 375 to 1500) were acquired in the Orbitrap with 120,000 resolution after accumulation of ions to a 4 × 10^5^ target value. Maximum injection time was set to 50 ms and dynamic exclusion was set to 30 s. MS2 analysis consisted of higher-energy collisional dissociation (HCD); AGC 1 × 10^5^; normalized collision energy (NCE) 38; maximum injection time 105 ms; 50,000 resolution and isolation window of 0.7 Da.

For proteome analysis, peptides were delivered to an analytical column and separated at a flow rate of 280 nL/min using a 105-min gradient from 5% to 30% of solvent B (solvent A, 0.1% formic acid (FA); solvent B, 0.1% FA and 99.9% acetonitrile).

### Phosphoproteome data preprocessing

Raw phosphoproteome MS data were processed using MaxQuant version 1.6.14.0 (Andromeda search engine). MS/MS spectra were searched against the UniProt human database with full tryptic specificity, up to two missed cleavage sites, carbamidomethylation of cysteine residues as a fixed modification, and phosphorylation of serine, threonine, and tyrosine, N-terminal protein acetylation, and methionine oxidation as variable modifications. Reporter ion intensities were processed by PSM-level ratio normalization. The false discovery rates for protein groups, peptide-spectrum matches, and phosphosites were set to < 0.01. PTM sites with localization probability > 0.75 were considered localized. Quantitative values of phosphorylation sites across fractions were automatically integrated and summarized in Phospho (STY) Sites.txt by MaxQuant. Peptides identified from the reversed database or annotated as potential contaminants were excluded from downstream analysis. Phosphosites were filtered and missing values were imputed using the same criteria and procedure described above for proteome data. Batch effects in phosphosite abundance were corrected using ComBat implemented in the R package sva version 3.58.0, with batch identity specified as the batch variable and a parametric empirical Bayes adjustment without additional covariates.

### Bulk RNA-seq sample preparation and preprocessing

MCF-7 cells were transfected with control siRNA (siControl) or siRNA targeting IκBα (siIκBα) at a final concentration of 25 nM and stimulated with TNFα (10 ng/mL) at the indicated time points. For bulk RNA-seq, siControl- and siIκBα-transfected cells were collected at 0, 3, 6, 12, and 48 h after TNFα stimulation. For washout experiments, TNFα was removed after 6 h of stimulation, and cells were collected at 12 or 48 h. Three biological replicates were prepared for each condition. Total RNA was isolated using the NucleoSpin RNA Plus kit (Macherey-Nagel; #740984.250), and library preparation and sequencing were performed by Rhelixa (Tokyo, Japan). Poly(A)+ RNA was isolated using the NEBNext Poly(A) mRNA Magnetic Isolation Module (New England Biolabs), and strand-specific libraries were prepared using the NEBNext Ultra II Directional RNA Library Prep Kit (New England Biolabs). Paired-end sequencing was performed on an Illumina NovaSeq X Plus platform. Raw sequencing reads were processed using the nf-core/rnaseq pipeline version 3.17.0. Raw sequencing reads were subjected to adapter and quality trimming using Trim Galore! version 0.6.10, followed by alignment to the GRCh38 genome using STAR version 2.7.11b. Transcript abundance was quantified using Salmon version 1.10.3, and gene-level expression counts were summarized based on the corresponding gene annotation.

### ATAC-seq data preprocessing

ATAC-seq data preprocessing was conducted as described previously^21^, and the processed data were reused in this study. Briefly, reads were aligned to GRCh38 using Bowtie2 version 2.3.4.3, and duplicate reads were removed using Picard version 2.20.7 and SAMtools version

1.9. Peaks were called using MACS2 version 2.2.4, filtered to exclude blacklist regions, and assessed for reproducibility using IDR version 2.0.4.2. Peak annotation and differential accessibility analysis were performed using HOMER version 4.9.1 and DESeq2 version 1.20.0, respectively.

### Single-cell (sc)RNA-seq sample preparation and preprocessing

MCF-7 cells were transfected with control siRNA (siControl) or siRNA targeting IκBα (siIκBα) at a final concentration of 25 nM and stimulated with TNFα (10 ng/mL) at the indicated time points. For single-cell RNA-seq, siControl- and siIκBα-transfected cells were collected at 0, 6, 12, and 48 h after TNFα stimulation. One sample was prepared for each condition. Cells were dissociated by trypsinization and prepared as single-cell suspensions. Single-cell RNA-seq libraries were generated by Rhelixa using the 10x Genomics single-cell 3’ RNA-seq platform with a GEM-X kit, with a target recovery of approximately 10,000 cells per sample. Libraries were sequenced on an Illumina NovaSeq X Plus platform. Raw sequencing data were processed using Cell Ranger version 9.0.1. Reads were aligned to the GRCh38 genome, and gene-by-cell count matrices were generated for each sample using the cellranger count pipeline with default parameters.

### Metabolome sample preparation and measurement

Metabolome analysis was performed as described previously^100^. Briefly, MCF-7 cell samples (approximately 1 × 10⁶ cells) were extracted with 1 mL of cold methanol containing 10- camphorsulfonic acid (1.5 nmol) and piperazine-1,4-bis(2-ethanesulfonic acid) (1.5 nmol) as internal standards. Samples were vortexed for 1 min, sonicated for 5 min, and centrifuged at 16,000 × g for 5 min at 4 °C. The supernatant (400 µL) was collected, and protein concentrations in the pellets were determined using a Pierce™ BCA Protein Assay Kit (Thermo Fisher Scientific, MA). The supernatant was mixed with 400 µL of chloroform and 320 µL of water, vortexed, and centrifuged at 16,000 × g for 5 min at 4 °C. The aqueous layer (500 µL) was collected, evaporated under vacuum, and stored at −80 °C until analysis. Dried extracts were reconstituted in 50 µL of water before analysis.

Hydrophilic metabolites were analyzed using two liquid chromatography high-resolution tandem mass spectrometry (LC-MS/MS) methods^100,101^. Anionic polar metabolites, including organic acids, sugar phosphates, and nucleotides, were analyzed using ion chromatography (Dionex ICS-5000+ HPIC system, Thermo Fisher Scientific) with a Dionex IonPac AS11-HC-4 μm column (2 mm i.d. × 250 mm, 4 μm particle size, Thermo Fisher Scientific) coupled with a Q Exactive high-performance benchtop quadrupole Orbitrap mass spectrometer (Thermo Fisher Scientific) (IC-MS/MS). Cationic polar metabolites, including amino acids, bases, and nucleosides, were analyzed using liquid chromatography (Nexera X2 UHPLC system, Shimadzu) with a Discovery HS F5 column (2.1 mm i.d. × 150 mm, 3 μm particle size, Merck) coupled with a Q Exactive instrument (PFPP-LC-MS/MS). The analytical platforms were operated using LabSolutions version 5.80 (Shimadzu) and Xcalibur version 4.2.47 (Thermo Fisher Scientific). Relative metabolite levels were calculated from peak areas normalized to the internal standard 10-camphorsulfonic acid and to the total protein amount of each sample.

### Integrated multi-omics analysis

TNF-induced responses were compared across transcriptome, proteome, and phosphoproteome datasets using values relative to the corresponding 0-h condition. RNA responses were represented as log₂ fold changes, whereas proteome and phosphoproteome responses were calculated as differences in preprocessed log₂ protein abundance and centered phosphosite abundance, respectively. Features with a Benjamini–Hochberg (BH)-adjusted P value < 0.05 and an absolute response ≥ 0.5 were considered significantly responsive. The numbers of responsive features were summarized at each time point for each omics layer.

Overall response burden was evaluated as the number of unique features showing a significant response at any time point and as the total number of significant feature–time point observations, and the IκBα KD-to-Control ratio was calculated for each measure.

TNF-response dynamics were further evaluated for features with complete trajectories through 48 h, with the response at 0 h defined as zero. For each feature, four descriptive measures were calculated: the absolute area under the response curve (AUC) from 0 to 48 h using trapezoidal integration, the maximum absolute response, the dynamic range defined as the difference between the maximum and minimum responses, and the maximum absolute slope between adjacent time points. These metrics were compared between Control and IκBα KD on a feature-by-feature basis within each omics layer.

### Kinase-substrate enrichment analysis

Kinase-substrate relationships were assigned using the human KINASE-PSP signatures in PTMsigDB version 2.0.0. Phosphosite sequence windows were normalized and matched to PTMsigDB based on sequences centered on the phosphorylation site. Kinases with at least five mapped phosphosites were retained for kinase-substrate enrichment analysis (KSEA).

For time-resolved KSEA, phosphosite abundance was averaged across three biological replicates at each time point, and responses were calculated relative to the corresponding 0-h value within each condition. Signed phosphosite responses were integrated using the trapezoidal rule over early (0–3 h) and late (12–48 h) time windows and divided by the respective window duration to obtain mean signed responses. For each kinase, the KSEA z- score was calculated from the mean response of its mapped substrates relative to the distribution of all measured phosphosites, normalized by the number of substrates. KSEA scores were interpreted as substrate-based estimates of kinase activity rather than direct measurements of kinase activation. One-tailed P values in the observed direction were calculated from the standard normal distribution and adjusted across kinases using the Benjamini–Hochberg (BH) method. To compare kinase responses between Control and IκBα KD, the signed perpendicular distance from the identity line was calculated as (z_IκBαKD − z_Control)/√2. Kinases with distances < −1 or > 1 were classified as Control-specific or IκBαKD-specific, respectively. Pearson and Spearman correlations between Control and IκBα KD KSEA z-scores were calculated for each time window using the R stats package version 4.5.2. For baseline KSEA, phosphosite levels at 0 h were compared between IκBα KD and Control cells using an empirical-Bayes linear model implemented in the R package limma version 3.66.0, with three biological replicates per condition. The resulting site-level IκBα KD- minus-Control differences were used directly for KSEA without significance-based prefiltering, using the same kinase-substrate mapping, minimum substrate number, and KSEA calculation described above.

### Metabolome data analysis

Metabolome data were analyzed using R. For each metabolite, differences between Control and IκBα KD, Control and TNF-treated Control, and Control and IκBα KD + TNF were assessed using unpaired two-sided Welch’s t tests assuming equal variance, with four biological replicates per condition. Metabolites with P < 0.05 were considered significantly altered and were used for subsequent pathway enrichment analysis. For pathway enrichment analysis, metabolites significantly altered in each comparison were identified using their Human Metabolome Database (HMDB) IDs and analyzed using the R package MetaboAnalystR version 4.3.0. Metabolites were cross-referenced by HMDB ID and mapped to the current Homo sapiens KEGG pathway library. Over-representation analysis was performed using the hypergeometric test, and P values were corrected for multiple testing using the BH method.

### Mitochondrial proteome data analysis

Organelle-associated protein sets, including mitochondrial proteins, were defined using Gene Ontology Cellular Component (GOCC) annotations with the R packages GO.db version 3.22.0 and org.Hs.eg.db version 3.22.0. At each time point, proteins were ranked according to the difference in mean abundance between IκBα KD and Control cells, and gene set enrichment analysis was performed using the R package fgsea version 1.36.2. Normalized enrichment scores (NES) and Benjamini–Hochberg-adjusted P values were used to evaluate enrichment. The top 30 leading-edge proteins from the mitochondrial gene set at 0 h were selected for heatmap visualization. Differences in protein abundance between IκBα KD and Control cells at each time point were evaluated using an empirical-Bayes linear model implemented in the R package limma version 3.66.0. Mitochondrial subcompartment annotations, including the outer membrane, inner membrane, matrix, and intermembrane space, were assigned based on GOCC. Protein changes with P < 0.05 and an absolute abundance difference ≥ 0.3 were highlighted in the heatmap. For mitochondrial targeting peptide analysis, reviewed human protein sequences obtained from UniProt were subjected to mTP score prediction using the TargetP 2.0 web server (https://services.healthtech.dtu.dk/services/TargetP-2.0/). Mitochondrial proteins were classified according to their mTP scores as low (mTP ≤ 0.2), intermediate (0.2 < mTP < 0.8), or high (mTP ≥ 0.8). At 0 h, protein abundance changes between IκBα KD and Control cells were compared between the low- and high-mTP groups using a two-sided Wilcoxon rank-sum test. High-mTP proteins at 0 h were further visualized by volcano plot, with P < 0.05 used to indicate significantly altered proteins.

### Integrated RNA-seq, proteomic, and ATAC-seq analysis and motif enrichment analysis

Baseline RNA-seq and proteome data were matched by gene symbol, retaining mitochondrial genes represented in both datasets. ATAC-seq peaks were linked to these genes using the existing annotations and restricted to regions within 100 kb of the annotated transcription start site. Multiple peaks associated with the same gene were retained as separate regions. Genes were classified as RNA-related (RNA log₂ fold change ≤ −0.3), protein-only (protein log₂ fold change ≤ −0.3 without meeting the RNA-related criterion), or reference (remaining genes).

Peak-level ATAC-seq log₂ fold changes were summarized as mean ± SE and compared between each reduction group and the reference group using a linear model with Dunnett- adjusted comparisons. For motif enrichment analysis, peaks with negative ATAC-seq log₂ fold changes in the RNA-related and protein-only groups were selected, with reference-group peaks used as controls. Peak-centered 200-bp sequences were extracted from hg38 and analyzed using AME version 5.5.9 with the JASPAR 2024 CORE vertebrate non-redundant database, Fisher’s exact test, and total-hit scoring. Motifs corresponding to PPARA, CREB1, YY1, RELA, and NFKB1 were visualized using nominal P values, retaining the motif with the smallest P value for each transcription factor and group.

### Single-cell RNA-seq data analysis

Single-cell RNA-seq data were analyzed using Seurat version 5.5.0 in R version 4.5.2. RNA assay layers in the merged Seurat object were joined before filtering. The mitochondrial count percentage was calculated using genes with symbols beginning with MT-. Cells were retained if they had more than 2,500 and fewer than 7,500 detected genes and mitochondrial counts below 3% of total counts. Genes were retained if they were detected in at least three retained cells and had a mean raw count greater than 0.05 across all retained cells, including zeros. After cell and gene filtering, cells were randomly downsampled within each sample (orig.ident) to the minimum number of cells remaining among samples, using a random seed of 319. Counts were normalized using Seurat LogNormalize with a scale factor of 10,000, and the normalized RNA data layer was used for downstream analysis. Cell-cycle effects were regressed out using S and G2/M scores. The 2,000 most variable genes were used for principal component analysis (PCA), and clustering was performed using the first 30 principal components with a resolution of 0.4. UMAP visualization was generated using the first 30 principal components with 30 nearest neighbors and a minimum distance of 0.3. Clusters were manually annotated as C0–C5 based on sample distribution. Differential expression between C5 and C3 was assessed using the Seurat FindMarkers function with the Wilcoxon rank-sum test, min.pct = 0.05, and logfc.threshold = 0.1. P values were adjusted using the Bonferroni method, and genes with adjusted P < 0.01 were considered differentially expressed. KEGG pathway enrichment analysis was performed using clusterProfiler version 4.18.4 and org.Hs.eg.db version 3.22.0. Gene symbols were converted to Entrez IDs, and all genes detected in the Seurat object were used as the background gene set. Pathways containing 10–500 genes were tested, and enrichment P values were adjusted using the Benjamini–Hochberg method. For visualization, pathways with nominal P < 0.01 were selected from the Cellular Processes and Metabolism categories, excluding disease-related pathways.

### RNA velocity and trajectory analysis

For RNA velocity analysis, spliced and unspliced count matrices generated using velocyto were integrated with the single-cell RNA-seq data, and RNA velocity was estimated using cell2fate version 0.1a0. For model training, up to 2,000 cells per cluster and 2,000 highly variable genes were selected without excluding any clusters. The Cell2fate_DynamicalModel was trained using spliced and unspliced counts for 200 epochs with a batch size of 512. Posterior estimates were obtained using 20 samples per cell. Trajectory structure was analyzed using scVelo version 0.2.4 and CellRank version 2.0.7. Velocity-directed partition-based graph abstraction (PAGA) was performed using scVelo with the six identified clusters (C0–C5), the existing nearest- neighbor graph, and the estimated RNA velocity. Cell fate probabilities were estimated using CellRank by combining a velocity kernel with a real-time kernel generated from experimental time information using moscot version 0.5.0 at a weight ratio of 0.6:0.4. C3 and C5 were defined as terminal states, and the 500 cells with the highest velocity pseudotime values in each terminal cluster were used to represent the terminal states. Fate probabilities toward C3 and C5 were calculated using the GPCCA estimator. Velocity pseudotime was normalized within each branch and used to visualize pseudotime-dependent gene expression patterns. To identify genes associated with C5 fate commitment, gene expression in C0 cells was correlated with C5 fate probabilities using Spearman correlation, followed by Benjamini–Hochberg correction. Genes ranked by the correlation coefficient were subjected to KEGG gene set enrichment analysis using clusterProfiler version 4.18.4.

### Trajectory inference using optimal transport

Optimal Transport (OT) was applied on the LogNormalized, filtered sc-RNAseq data set, considering the top 2,000 highly variable genes. Individual transport maps were calculated for the transitions from 0–6 h, 6–12 h and 12–48 h using the Python library Waddington OT^102^ version 1.0.8. In the applied unbalanced OT, an initial estimate on the expected increase in mass is required. This estimate of cell specific birth-death-rates was calculated by a logistic scaling of gene-set expression scores of proliferation and apoptosis related genes derived from Tirosh et al.^103^ and Liberzon et al.^104^. The cell specific birth-death-rates were globally adjusted to match experimentally derived bulk counts of cells for each time-point. These bulk-calibrated rates were then used to solve unbalanced optimal transport problem with parameters, λ_1_ = 25, λ_2_ = 50 and *ε* = 0.05, where λ_1,_ λ_2_ and *ε* are the coefficients for the relaxation terms in the optimization. The resulting cell-by-cell transport map was aggregated according to the previously determined clusters (C0–C5) and normalized by the total transported mass.

### Apoptosis-related module score analysis

Apoptosis-related transcriptional activity was evaluated using the Seurat AddModuleScore function. The pro-apoptotic gene set comprised BCL2L11, BBC3, PMAIP1, BAX, BAK1, BID, BAD, and BIK, whereas the anti-apoptotic gene set comprised BCL2, BCL2L1, BCL2L2, MCL1, and BCL2A1. For statistical comparisons, 50 cells were randomly selected from each cluster (C3 and C5), scores were compared between C3 and C5 using the Wilcoxon rank-sum test with Benjamini–Hochberg correction.

## Notes

### Competing Interest Statement

The authors have declared no competing interest.

