## Supplementary Information for "IκBα loss impairs mitochondrial homeostasis and primes cells for senescence"

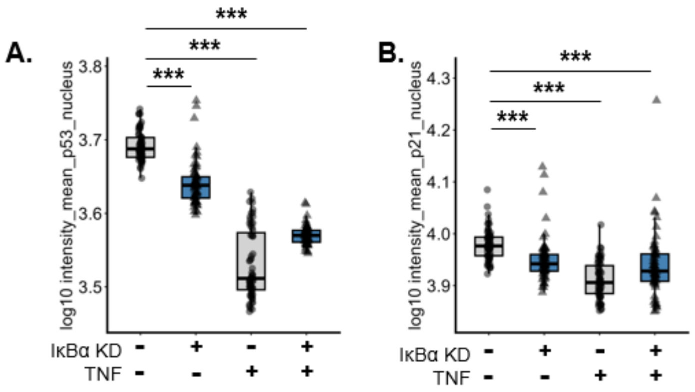


**Figure S1. Nuclear p53 and p21 levels following IκBα depletion and TNFα stimulation.** (A, B) Quantification of nuclear p53 (A) and p21 (B) fluorescence intensity in MCF-7 cells transfected with control or IκBα siRNA (25 nM) and cultured with or without TNFα (10 ng/mL) for 24 h. Cells were fixed and stained for p53 and p21 by 4i, with SYTOX Green counterstaining of nuclei. For each cell, the mean nuclear fluorescence intensity was measured, and values were averaged within each image (well × field). Each point represents one image. Values were $\log_{10}$-transformed, and box plots show the median and interquartile range. Statistical significance was determined using Dunnett's multiple comparisons test, with untreated control cells as the reference group. ***$P < 0.001$.

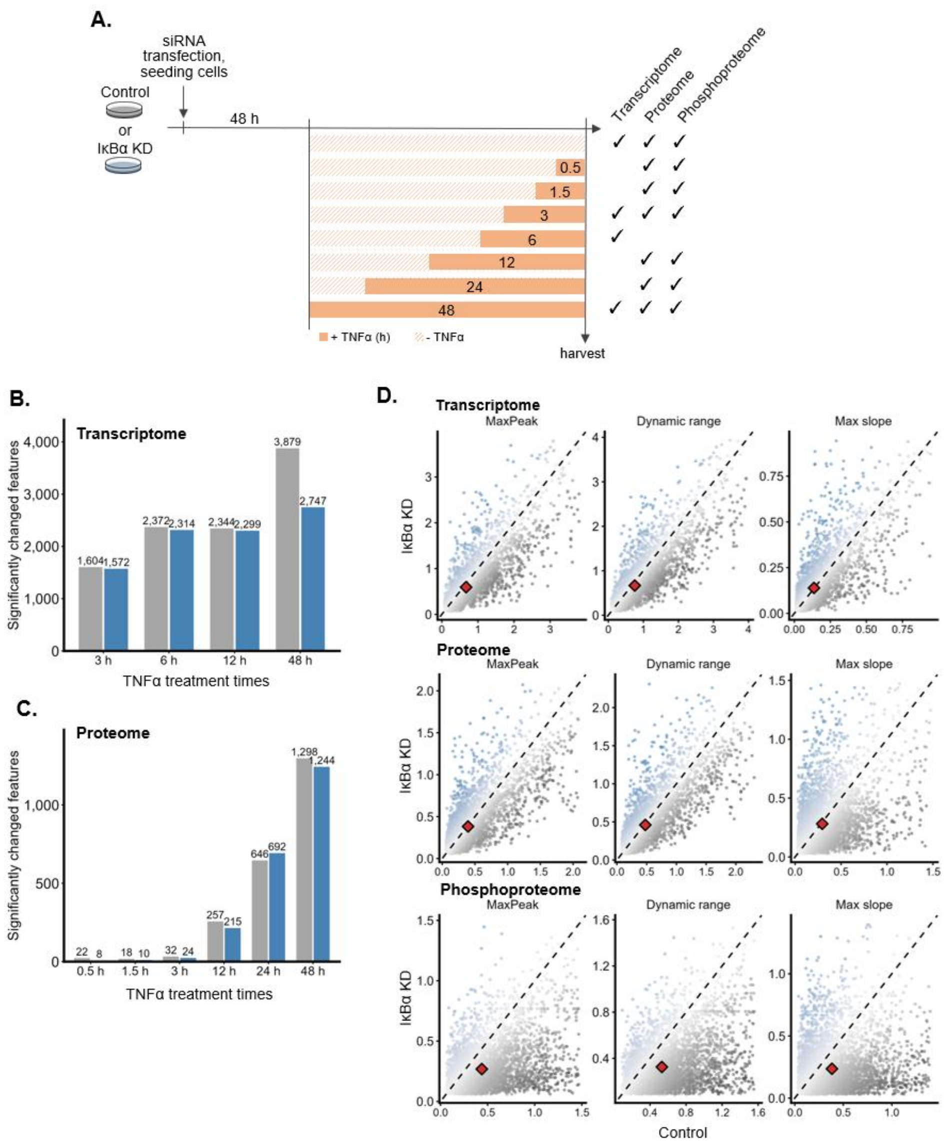


**Figure S2. Multi-omics profiling of TNFα response dynamics following IκBα depletion**

(A) Schematic overview of the experimental design and sampling time points for transcriptomic, proteomic, and phosphoproteomic analyses. Control and IκBα KD MCF-7 cells were stimulated with TNFα (10 ng/mL) 48 h after siRNA transfection. Samples were collected at the indicated time points for each omics analysis. (B) Number of TNF-responsive Genes at each time point in Control and IκBα KD cells. Significant TNF responses were defined as BH FDR < 0.05 and absolute response ≥ 0.5. (C) Number of TNF-responsive Proteins at each time point in Control and IκBα KD cells. Significant TNF responses were defined as BH FDR < 0.05 and absolute response ≥ 0.5. (D) MaxPeak, dynamic range, and maximum slope of TNFα responses in Control versus IκBα KD for features with complete 0–48 h trajectories. TNFα responses were calculated relative to the respective 0-h condition, with the 0-h response set to 0. Dashed line, y

= x; diamond, centroid. Up to 6,000 features per layer were randomly sampled and displayed within the 1st–99th percentile range.

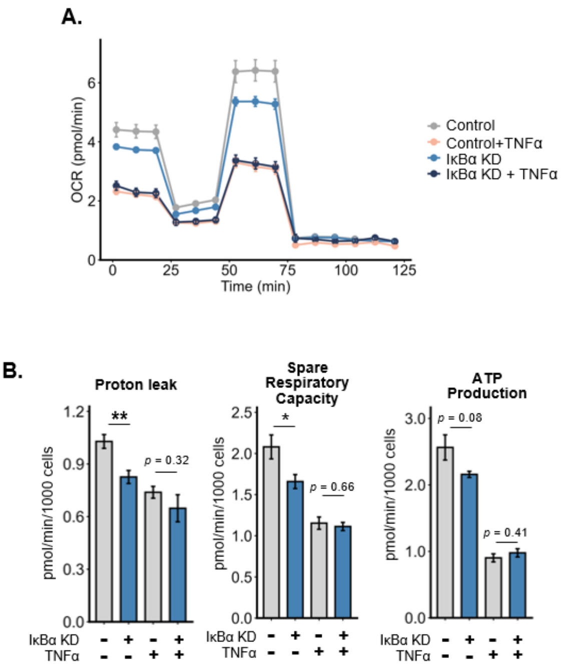


**Figure S3. IκBα depletion impairs mitochondrial respiration**

(A) Oxygen consumption rate (OCR) profiles of Control and IκBα KD MCF-7 cells with or without TNFα treatment (10 ng/mL, 48 h), measured by Seahorse analysis and normalized to cell number. Data are presented as mean ± SEM. (B) Proton leak (left), Spare respiratory capacity (middle), and ATP production (right) in Control and IκBα KD MCF-7 cells with or without TNFα treatment (10 ng/mL, 48 h), measured by Seahorse OCR analysis and normalized to cell number. Bars represent mean ± SEM. P values were determined using unpaired two-tailed Welch's t tests for the indicated comparisons.

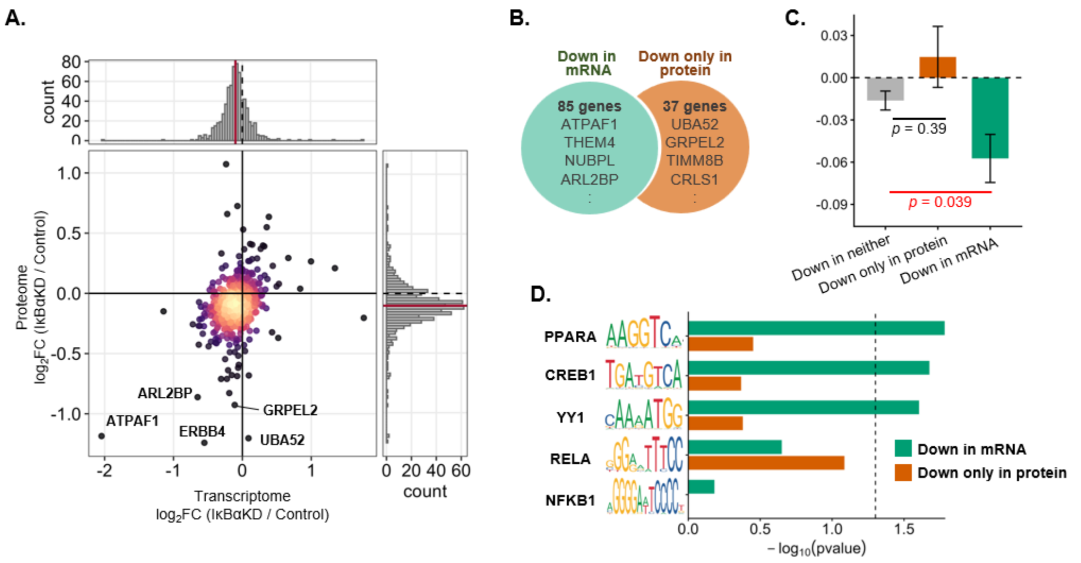


**Figure S4. Integrated analysis of mitochondrial gene expression, protein abundance, and chromatin accessibility following IκBα depletion**

(A) Comparison of RNA and protein abundance changes in mitochondrial genes following IκBα KD at 0 h. RNA-seq and proteome log2 fold changes are shown for genes annotated as mitochondrial by Gene Ontology Cellular Component (GOCC). Point color indicates local density. Marginal histograms show the corresponding distributions, with red lines indicating the mean. (B) Classification of mitochondrial genes based on RNA and protein changes. Genes with RNA-seq log2FC ≤ −0.3 were classified as Down in mRNA; genes not meeting this criterion but with proteome log2FC ≤ −0.3 were classified as Down only in protein; all remaining genes were classified as Down in neither. (C) ATAC-seq changes in the gene groups defined in (B). ATAC-seq peaks within ±100 kb of the transcription start site were included. Bars show mean log2FC ± SE. P values were determined by Dunnett's multiple-comparison test using Down in neither as the reference. (D) Transcription factor motif enrichment analysis of ATAC-seq peaks associated with the gene groups defined in (B). For Down in mRNA and Down only in protein, peaks with ATAC-seq log2FC < 0 were analyzed against all peaks from Down in neither. Sequences of 200 bp centered on each peak were analyzed by AME using the JASPAR 2024 CORE vertebrate motif database and Fisher's exact test. For each indicated transcription factor, the motif with the lowest P value is shown. The dashed line indicates P = 0.05.

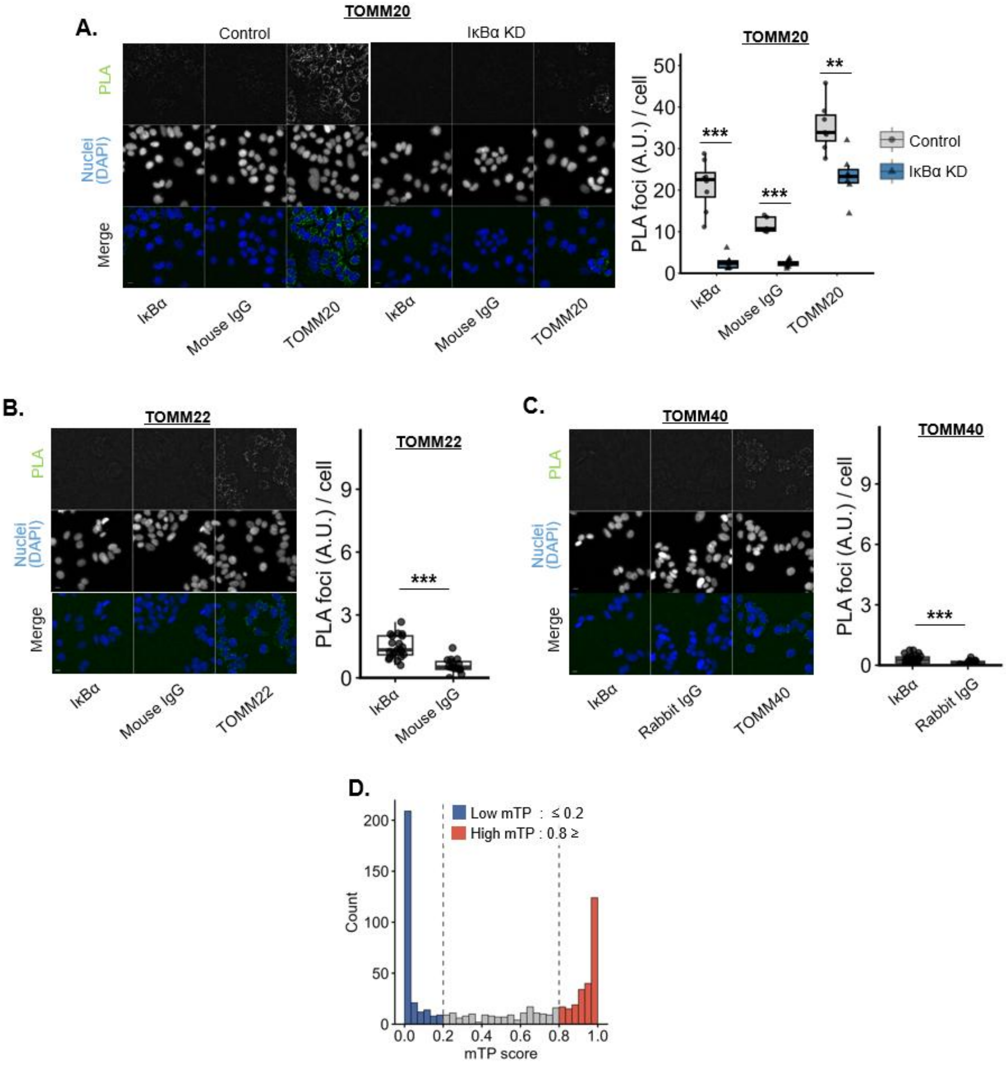


**Figure S5. Proximity ligation analysis of IκBα and TOM complex components**

(A) Proximity ligation assay (PLA) of IκBα and TOMM20 in MCF-7 cells. Left: Representative PLA images. Right: Quantification of PLA foci per cell in Control and IκBα KD cells using the indicated antibodies. (B) Proximity ligation assay (PLA) of IκBα and TOMM22 in MCF-7 cells. Left: Representative PLA images. Right: Quantification of PLA foci per cell compared with the Mouse IgG negative control. (C) Proximity ligation assay (PLA) of IκBα and TOMM40 in MCF-7 cells. Left: Representative PLA images. Right: Quantification of PLA foci per cell compared with the Rabbit IgG negative control. (D) Histogram of mTP scores for 739 mitochondria-related genes. Each point represents the mean value per image (well × field). Statistical significance was assessed using two-sided Welch's t-tests. In (A) and (B), comparisons were made against the corresponding IgG negative controls; in (C), Control and IκBα KD were compared within each antibody condition.

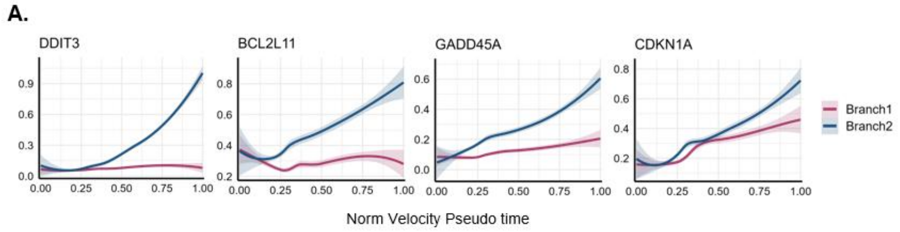


**Figure S6. Branch-specific expression dynamics of apoptosis-associated genes and CDKN1A**

(A) Velocity pseudotime expression profiles of apoptosis-related genes (DDIT3, BCL2L11, and GADD45A) and the senescence-related gene CDKN1A. Lines indicate fitted curves, and shaded areas indicate 95% confidence intervals. Velocity pseudotime was scaled from 0 to 1 based on the minimum and maximum values within each branch.

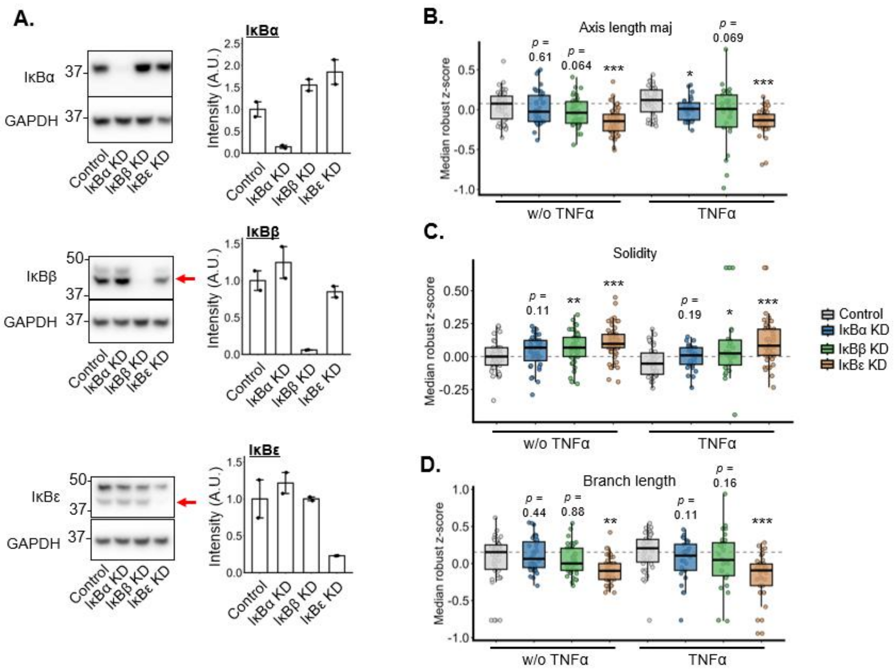


**Figure S7. Validation of IκB isoform knockdown and quantification of mitochondrial morphology**

(A) Western blot analysis of IκBα, IκBβ, and IκBε in MCF-7 cells transfected with negative-control siRNA or siRNAs targeting each IκB isoform (25 nM). Left: Representative immunoblots, with GAPDH as a loading control. Red arrows indicate the target bands. Right: Quantification of protein levels normalized to the corresponding control condition. Bars represent mean ± SE, and dots indicate individual replicates (n = 2). (B) Field-level median robust z-scores for mitochondrial branch length in Control and IκBα, IκBβ, or IκBε KD cells with or without TNF treatment. (C) Field-level median robust z-scores for mitochondrial major-axis length.

(D) Field-level median robust z-scores for mitochondrial solidity. For (B)–(D), each point represents one field, and box plots show the median and interquartile range. Dashed horizontal lines indicate the median value of untreated Control fields. n = 28–54 fields of view per group. P values in (B)–(D) were calculated using two-sided Welch's t-tests comparing each IκB KD group with the corresponding Control group within each treatment condition, followed by Benjamini–Hochberg correction for multiple comparisons. *P < 0.05, **P < 0.01, ***P < 0.001.

A.

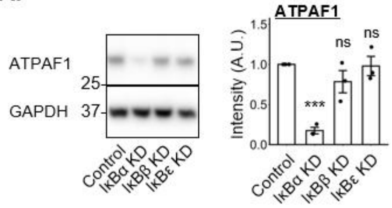


**Figure S8. IκBα depletion selectively reduces ATPAF1 protein levels among canonical IκB isoforms**

(A) Western blot analysis of ATPAF1 in MCF-7 cells transfected with negative-control siRNA or siRNAs targeting IκBα, IκBβ, or IκBε (25 nM). Left: Representative immunoblots, with GAPDH as a loading control. Right: Quantification of ATPAF1 protein levels normalized to the control condition. Bars represent mean ± SE, and dots indicate individual replicates (n = 3). P values were determined separately under untreated and TNFα-treated conditions using Dunnett's multiple comparisons test, with each KD condition compared with the corresponding Control. *P < 0.05, **P < 0.01, ***P < 0.001, ****P < 0.0001; ns, not significant.

### SUPPLEMENTAL INFORMATION

#### Table 1

| **Oligonucleotides** | | |
|---|---|---|
| REAGENT or RESOURCE | SOURCE | IDENTIFIER |
| qRT-PCR forward primer: RPL27 5'-CTGTCGTCAATAAGGATGTCT-3' | This paper | N/A |
| qRT-PCR reverse primer: RPL27 5'-CTTGTTCTTGCCTGTCTTGT-3' | This paper | N/A |
| qRT-PCR forward primer: IL-6 5'-CCAGAGCTGTGCAGATGAGT-3' | This paper | N/A |
| qRT-PCR reverse primer: IL-6 5'-TTTCTGCAGGAACTGGATCAGG-3' | This paper | N/A |
| qRT-PCR forward primer: IL-8 5'-CCTGATTTCTGCAGCTCTGTGT-3' | This paper | N/A |
| qRT-PCR reverse primer: IL-8 5'-GGTGGAAAGGTTTGGAGTATGTCT-3' | This paper | N/A |
| qRT-PCR forward primer: CCL2 5'-AAGACCATTGTGGCCAAGGA-3' | This paper | N/A |
| qRT-PCR reverse primer: CCL2 5'-TTCGGAGTTTGGGTTTGCT-3' | This paper | N/A |
